# A wheat resistosome defines common principles of immune receptor channels

**DOI:** 10.1101/2022.03.23.485489

**Authors:** Alexander Förderer, Ertong Li, Aaron Lawson, Ya-nan Deng, Yue Sun, Elke Logemann, Xiaoxiao Zhang, Jie Wen, Zhifu Han, Junbiao Chang, Yuhang Chen, Paul Schulze-Lefert, Jijie Chai

## Abstract

Plant intracellular nucleotide-binding leucine-rich repeat (NLRs) receptors detect pathogen effectors to trigger immune responses. Indirect recognition of a pathogen effector by the dicotyledonous *Arabidopsis thaliana* coiled-coil (CC) domain containing NLR (CNL) ZAR1 induces the formation of a large hetero-oligomeric protein complex, termed the ZAR1 resistosome, which functions as a calcium channel required for ZAR1-mediated immunity (*1*–*3*). Whether the resistosome and channel activities are conserved among plant CNLs remains unknown. We report here a cryogenic electron microscopy (cryo-EM) structure of the wheat CNL Sr35 in complex with the effector AvrSr35 of the wheat stem rust pathogen at 3.0 Å resolution. Direct effector binding to the leucine-rich repeats (LRRs) of Sr35 results in the formation of a pentameric Sr35-AvrSr35 complex, which we designate the Sr35 resistosome. Wheat Sr35 and Arabidopsis ZAR1 resistosomes bear striking structural similarity, including a previously unnoticed arginine cluster in the LRR domain that co-occurs and forms intramolecular interactions with the ‘EDVID’ motif in the CC domain. Electrophysiological measurements show that the Sr35 resistosome exhibits non-selective cation channel activity. These structural insights allowed us to generate novel variants of closely related wheat and barley orphan NLRs that recognize AvrSr35. Our data support the evolutionary conservation of CNL resistosomes in plants and demonstrate proof of principle for structure-based engineering of NLRs for crop improvement.

## INTRODUCTION

Plant NLRs are intracellular receptors that play a key role in the plant innate immune system by sensing the presence of pathogen effectors delivered inside plant cells during pathogenesis through direct or indirect recognition (*4*, *5*). Activation of plant NLRs generally leads to an array of immune responses, often linked to rapid host cell death at sites of attempted pathogen infection. Structural and functional homologs of plant NLRs evolved from independent events for intracellular non-self perception in animal innate immunity and are characterized by their conserved central nucleotide-binding and oligomerization domains (NOD) (*6*). Plant NLRs can be broadly separated into two classes: those with an N-terminal CC domain or an N-terminal Toll/interleukin 1 receptor (TIR) domain. Among the flowering plants dicots typically possess both receptor classes, whilst monocots, including cereals, only encode CNL receptors (*7*).

Wheat stem rust caused by fungal infection with *Puccinia graminis* f sp. *tritici (Pgt*) threatens global wheat production (*8*), and the emergence of widely virulent *Pgt* strains, including the Ug99 lineage, has motivated the search for stem rust resistance in wheat germplasm and wild relatives over the past two decades. This resulted in the isolation of eight stem rust resistance (*Sr*) genes that are phylogenetically related and belong to a clade of grass CNLs all of which confer strain-specific immunity (*9*–*15*) (‘clade I’ CNLs defined by ref. (*16*)). The MLA receptors of the wheat sister species barley belong to the same group as the latter receptors and share strain-specific immunity with *Sr* genes (*16*). *Sr35* was first identified in a *Triticum monococcum* (Einkorn) landrace, is lacking in hexaploid bread wheat (*T. aestivum*)and its A-genome diploid donor *T. urartu,* but confers immunity to *Pgt* Ug99 in bread wheat when transferred as a transgene (*14*). *Sr35* resistance has been linked to the recognition of the *Pgt* effector *AvrSr35* (*17*), but until now, it has remained inconclusive whether Sr35 receptor-mediated host cell death is driven by direct physical interaction with the AvrSr35 effector (*17*, *18*).

## RESULTS

### Purification and cryo-EM reconstruction of the Sr35 resistosome

To purify Sr35, we expressed the protein alone or together with AvrSr35 in Sf21 insect cells. Unexpectedly, cell death was observed when the receptor was co-expressed with *AvrSr35* (Fig. 1A), suggesting that Sr35 and its effector are sufficient to mediate this immunity-associated response in insect cells in the absence of other plant proteins. To circumvent cell death activation for protein purification, we introduced substitutions in the N-terminal residues L15E/L19E (Sr35^L15E/L19E^), which are predicted to be essential for Sr35 membrane association by analogy with the ZAR1 resistosome (*2*). Mutational analysis of the corresponding N-terminal residues of the tomato CNL NRC4 have been shown to abrogate cell death activity in *Nicotiana benthamiana* (*19*). Indeed, expression of the *Sr35^L15E/L19E^* variant markedly reduced *Sr35*-induced cell death in insect cells (Fig. 1A). Using affinity-tagged Sr35^L15E/L19E^, co-expressed with affinity-tagged AvrSr35, we were able to enrich the Sr35-AvrSr35 complex for subsequent separation of potential receptor complex isoforms and correctly folded receptor complexes by size-exclusion chromatography (SEC; fig. S1B, C). In SEC, the affinity-purified protein complex eluted with a broad peak with a maximum UV absorbance of approximately 550 kDa (70 mL). Individual fractions were analysed *via* negative staining and a large number of star-shaped particles with five-fold symmetry were identified (fractions at 60–69 mL mL; Fig. 1A). The most monodisperse fractions were pooled and used for cryo-EM analysis.

**FIG. 1.**
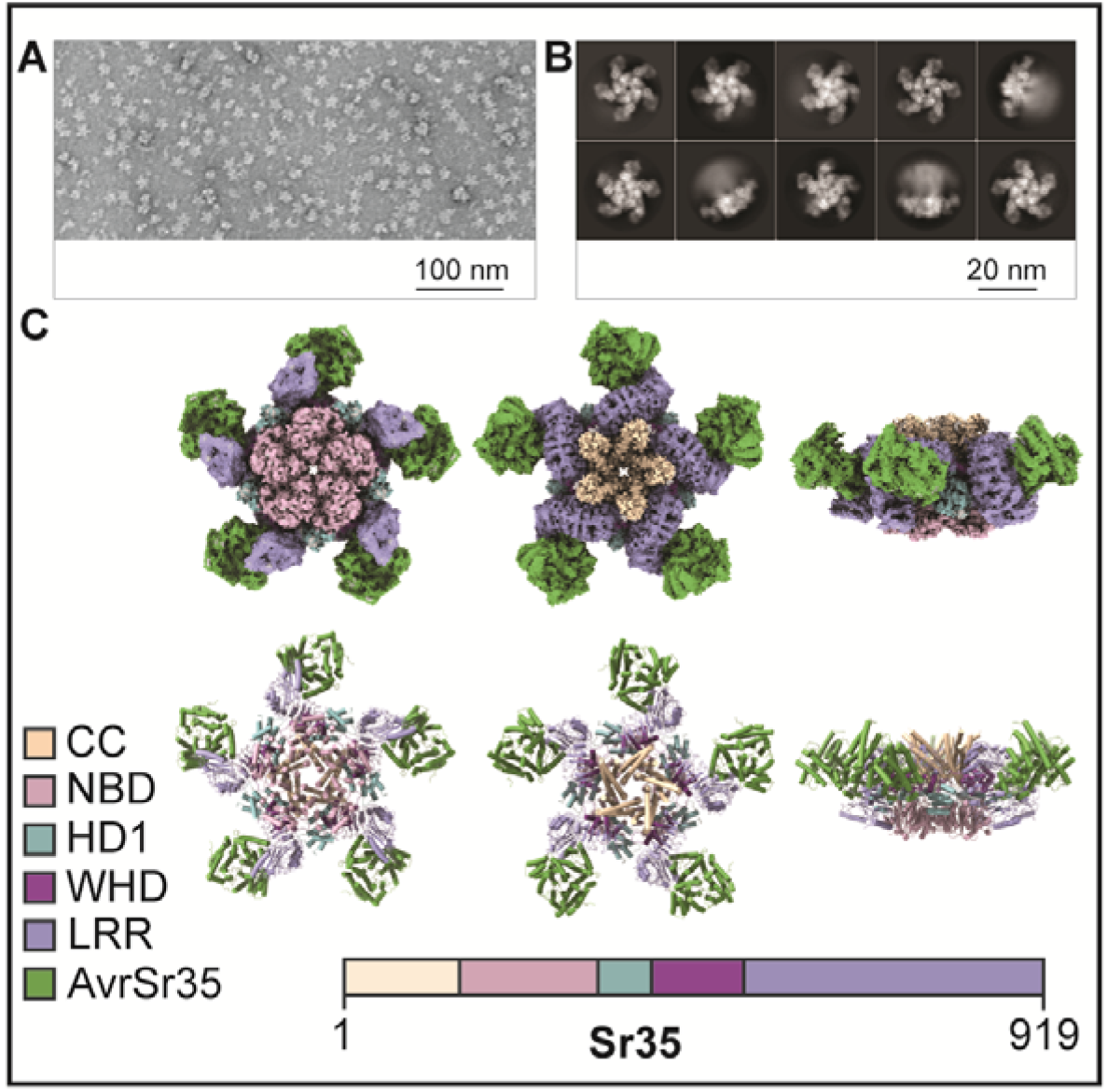
3D reconstruction of Sr35 resistosome (**A**) Negative staining of purified Sr35 in complex with AvrSr35 (Sr35 resistosome). Star-shaped particles were enriched by affinity purification and size-exclusion chromatography. Monodisperse Sr35 resistosomes average in size with approximately 24 nm. (**B**) 2D classifications of the Sr35 resistosomes from cryo-EM sample. Particles show preferential orientation for bottom and top view. Fewer but sufficient particles are in side view. (**C**) (top) Cryo-EM density map with 3 A and the finally refined structure model (bottom) of the Sr35 resistosome shown in three different orientations. AvrSr35 is coloured green and Sr35 domains are shown according to the colour code shown in the figure legend. From left to right bottom, top and side view.

We analysed the Sr35-AvrSr35 complex sample by cryo-EM (refer to fig. S2 for analysis flowchart) using a total of 1,608,441 individual particles for reference-free two-dimensional (2D) classification (Fig. 1B). After three-dimensional (3D) classification, a subset of 230,485 particles was used for reconstruction, yielding a density map of 3.0 Å (Fig. 1C, top). Despite the high resolution of up to 2.5 Å in the centre of the complex, the local resolution decreased towards the outer edge to approximately 4 Å (fig. S2C), indicating that the outer region of the complex is more flexible. To compensate for this decreased resolution, a local mask was used for the outer region, yielding a local density map with a resolution of 3.33 Å (fig. S2C). Both density maps were used for model building (Fig. 1C, bottom).

The final 3D reconstruction of the Sr35-AvrSr35 complex contains five receptor protomers each bound to one effector molecule. The reconstruction revealed a wheel-like structure, of a similar size to the ZAR1 resistosome (*2*), that we termed Sr35 resistosome. As in the ZAR1 resistosome, five Sr35 NOD modules define the base of the circular protomer arrangement, and a helical barrel formed by the five CC domains is buried at the centre. Unlike ZAR1, the LRR domains at the outer region do not pack against each other in the Sr35 resistosome, which might explain why this region is more flexible. AvrSr35 adopts an exclusively helical fold (fig. S3). A 3D structure search using DALI (*20*) showed that there are no other known proteins structures sharing the AvrSr35 fold. Five AvrSr35 proteins bind exclusively to the C-terminal part of the LRR domains in the complex.

### Sr35 oligomerization and stabilization features are evolutionarily conserved

In plants, the central NOD module of NLRs is subdivided into a nucleotide-binding domain (NBD), helical domain 1 (HD1) and winged helical domain (WHD). ATP/dATP has been shown to be important for ZAR1 oligomerization as it stabilizes the active conformation of ZAR1 via its interaction with the WHD in the NOD module. There is an unambiguous cryo-EM density at the predicted nucleotide-binding site between the HD1 and NBD domains that is unfilled by Sr35 and AvrSr35. The ATP molecule fits well into this cryo-EM density. The modelled ATP is nested in a groove formed by HD1 and NBD. The short α-helix that mediates inter-protomer interaction (Fig. 2A and C) also caps the ATP molecule. In contrast to ZAR1, ATP does not directly contact the WHD of Sr35. Instead, the γ-phosphate group of ATP forms a bidentate hydrogen bond with Sr35 NBD^R157^ and NBD^R311^ (Fig. 2C). The latter also forms a hydrogen bond with Sr35 WHD^S420^ (Fig. 2C), showing an indirect coupling of the ATP γ-phosphate group with the WHD of Sr35.

**FIG. 2.**
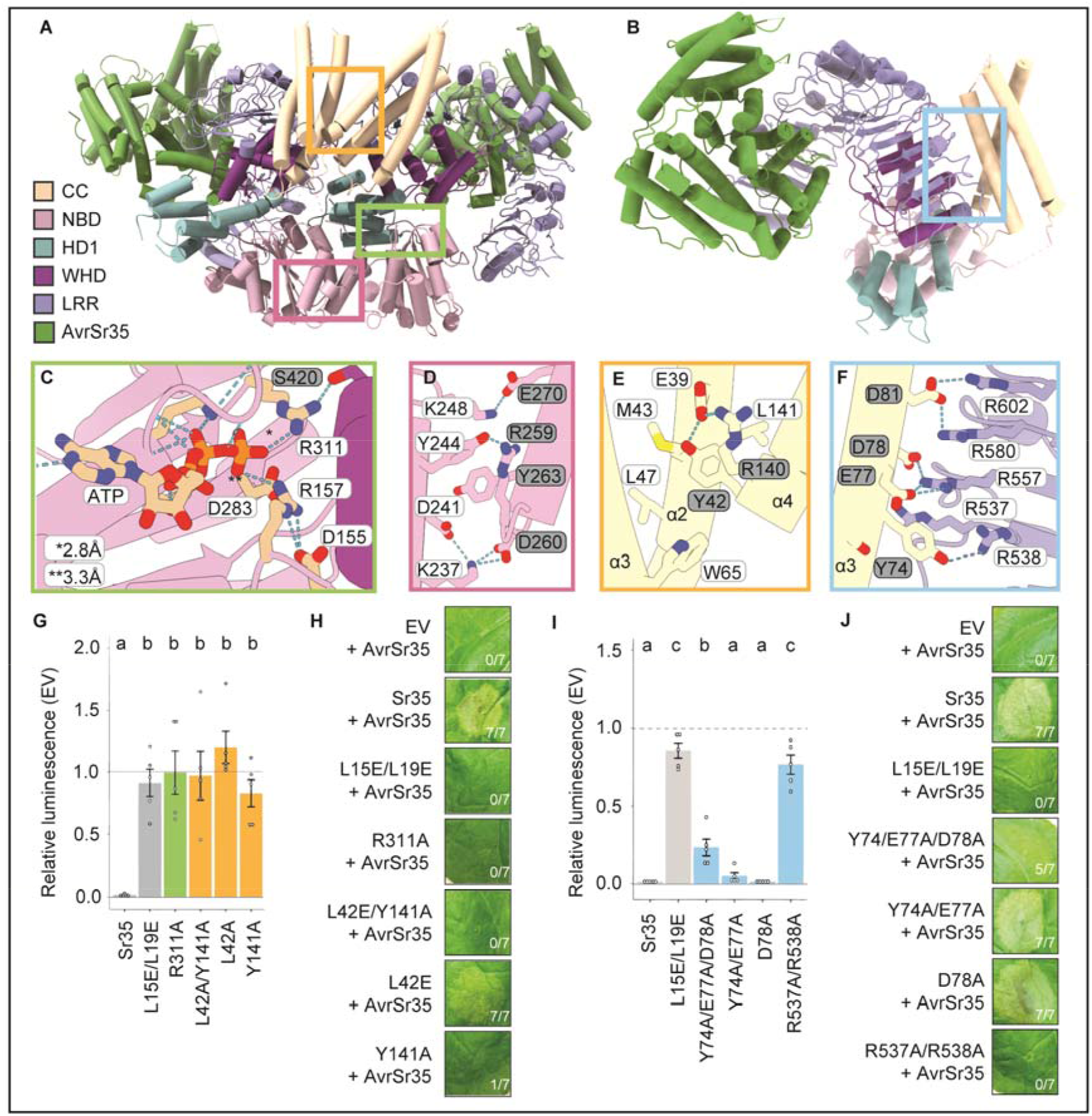
Critical aspects for the assembly of theSr35 resistosome (**A**) Sr35 resistosome structural model showing a lateral dimer used to design point mutations. Boxes in green, yellow and pink indicate positions of the zoomed views of (C to F). Sr35 domains and AvrSr35 are coloured according to in-figure legend. Box in blue indicating relative position of structural detail shown in (F). (**B**) Structure of one Sr35 protomer in complex with AvrSr35. Colour code as in (A). (**C**) Structural detail of the ATP binding pocket of one protomer bound to an ATP molecule. Note the specific hydrogen bond of R311 with gamma-phosphate group of ATP at a 2.8 Å distance. R311 was subjected to mutational analysis in (G to H). Grey and white residue label boxes corresponding to NBD and WHD residues respectively. (**D**) Structural detail of the interface between the NBDs of a lateral dimer. Dashed lines represent polar interactions. Grey and white residue label boxes corresponding to two neighbouring protomers from the pentamer. (**E**) Structural detail of the interface between the CC domains of a lateral dimer. (**F**) Structural detail of CC and LRR domain interface within one Sr35 protomer. Acidic residues in the EDIVD motif of the CC domain form salt bridges with positive Arg (R) residues of the R-cluster in the LRR domain. (**G**) Co-transfection of *Sr35* and *Sr35* mutants with *AvrS35* in wheat protoplasts. Relative luminescence was used as a readout for cell death. An empty vector treatment was used as the relative luminescence baseline of 1 (mean ±SEM; *n* = 5). Statistical analysis with one-way ANOVA and Tukey’s test. Treatments labelled with different letters differed significantly (*p* < 0.05). Additional statistical values reported in methods. Bar colours correspond to box colours in (C and D). (**H**) Tobacco cell death data of *Sr35* and *Sr35* mutants with *AvrSr35*. Representative data is shown from 7 replicates and scored for leaf cell death. (**I**) Wheat protoplast data of EDIVD and R-cluster mutants. Experiment performed as in (G). Bar colours correspond to box colour in F. (**J**) *N. benthamiana* cell death data of EDVID and R-cluster mutations. Representative data is shown from 7 replicates and scored for leaf cell death.

Similar to the ZAR1 resistosome, NBD-NBD contacts contribute to Sr35 protomer packing (Fig. 2D). Sr35 NBD^Y244^ from one protomer packs tightly against Sr35 NBD^R259^ and Sr35 NBD^Y263^ from an adjacent protomer (Fig. 2D). Additionally, a hydrogen bond is established between Sr35 NBD^Y244^ and Sr35 NBD^R259^. Of note, the CC domain of Sr35 contributes considerably to the inter-protomer interactions (Fig. 2A): the C-terminal half of α4-helix from one protomer packs against the C-terminal sides of α2- and α4-helices of the neighbouring CC protomer. At the centre of this interface is CC^Y141^, which makes extensive hydrophobic contacts with Sr35 CC^L42^, CC^M43^, CC^L47^ and CC^W65^ (Fig. 2E). Moreover, CC^Y141^ participates in a hydrogen bonding triad together with CC^R140^ from the same and CC^E39^ from the neighbouring protomer (Fig. 2E). As previously reported (*21*), the long linker region between the CC and NBD domain is also involved in mediating oligomerization of the Sr35 resistosome.

To functionally test the requirements for these interactions in mediating the assembly of the Sr35 hetero-oligomeric complex, we introduced amino acid substitutions into the receptor and assessed their impact on *Sr35*-mediated cell death using a luciferase (LUC) activity assay in wheat protoplasts (*22*) prepared from a genotype that does not recognize AvrSr35 (cultivar ‘Chinese Spring’). In this protoplast transfection assay, the relative (to empty vector, EV) luminescence of the *LUC* reporter is an indicator of cell viability. Co-transfection of *Sr35, AvrSr35* and the *LUC* reporter resulted in a near complete loss of luminescence signal, indicating massive cell death of the protoplasts and suggesting receptor activation by AvrSr35 (Fig. 2G). Consistent with the insect cell data described above, wheat protoplasts co-expressing *Sr35^L15E/L19E^* and *AvrSr35* displayed luminescence levels that were comparable to those co-expressing *EV* and *AvrSr35* constructs, indicating that the cell death activity of the Sr35^L15E/L19E^ receptor is substantially impaired (Fig. 2G). A similarly drastic loss of receptor-mediated cell death activity was observed with substitutions predicted to affect CC inter-protomer interactions (Y141A, L42E and L42E/Y141A) or the ATP-binding site (R311A) (Fig. 2G).

To corroborate the data from wheat protoplasts, we employed *Agrobacterium tumefaciens*-mediated transient gene expression of *Sr35* and *AvrSr35* in *N. benthamiana* leaves. Co-expression of *Sr35* and *AvrSr35*, but not *AvrSr35* plus EV, resulted in cell death in the *Agrobacterium-infiltrated* area (Fig. 2H). In contrast, cell death was abolished when *AvrSr35* was co-expressed with the *Sr35* mutants that are predicted to perturb Sr35 oligomerization (Fig. 2H), with the exception of Sr35^L42E^, which showed residual cell death activity only in *N. benthamiana. In planta*, protein levels of wild-type Sr35 and all receptor mutants tested were comparable, indicating that these substitutions do not render the receptor unstable (fig. S4A). Together, these data strongly suggest that the residues mediating Sr35 oligomerization in the cryo-EM structure are necessary for cell death activity in wheat and heterologous *N. benthamiana*.

A conserved sequence in the CC domain, long known as the ‘EDVID (Glu-Asp-Val-Ile-Asp) motif’ that is present in approximately 38% of Arabidopsis CNLs (*23*) and first described in the potato CNL Rx, is used to group CNLs with or without this motif, but its function has remained unclear (*23*, *24*)). In the cryo-EM structure of the Sr35 resistosome, the ED<u>IV</u>D motif (Glu-Asp-<u>Ile-Val</u>-Asp) and the adjacent Y^74^ mediate the packing of the LRR domain against the CC domain. Acidic residues from the motif form strong contacts with five arginine residues in the LRR domain (LRR^R537^, LRR^R538^, LRR^R557^, LRR^R580^, and LRR^R602^). These contacts comprise two bidentate salt bonds and a cation-π interaction (Fig. 2B and F). The extensive contacts in this region are further reinforced by hydrogen bonding and van der Waals contacts. Interestingly, these arginine residues are separated from each other by LRR repeats, which results in their spatial separation along the primary amino acid sequence (fig. S5A). Re-inspection of the ZAR1 resistosome shows that similar intramolecular interactions exist between arginine residues in the LRR and ‘EDIVD’ of Sr35. In both resistosomes the respective arginine residues cluster together and form a positively charged surface patch (fig. S5B). We therefore term this resistosome region LRR^R-cluster^. Separation of the arginine residues by a full leucine-rich repeat along the primary sequence explains why the LRR^R-cluster^ had remained unnoticed. A sequence alignment of CNLs shows that the LRR^R-cluster^ is conserved and co-occurs with the EDVID motif (fig. S5A).

To test whether the LRR^R-cluster^ is important for *Sr35*-mediated cell death, we substituted residues from the interface between the arginine cluster and the EDVID motif and assessed the impact of these mutations on cell death activity using the wheat protoplast and *N. benthamiana* leaf assays described above. Simultaneous mutations of LRR^R537A/R538A^ in the LRR^R-cluster^ essentially abolished cell death activity (Fig. 2H and I). Similarly, a triple substitution in the Sr35 EDVID motif, including the adjacent Y^74^, (Y74A/E77A/D78A) reduced or abolished cell death activity in protoplasts and *N. benthamiana*, respectively, without affecting NLR stability (fig. S4B). These observations suggest that the co-occurrence of the EDVID motif and LRR^R-cluster^ is an evolutionarily conserved stabilization mechanism of CNL resistosomes. As the EDVID-LRR^R-cluster^ interactions are also present in the inactive ZAR1 and Alphafold 2-modelled Sr35 monomers and an extensive fold switching occurs in the CC domain during receptor activation (fig. S5C), these intramolecular interactions must be transiently disrupted to allow α1-helix to flip.

### The Sr35 resistosome has nonselective calcium channel activity

Albeit they share only 28.4% sequence conservation and although the funnel-shaped structure composed of the α1-helix in the ZAR1 resistosome is not well defined in the Sr35 resistosome, the structures of the wheat and Arabidopsis ZAR1 resistosomes are highly similar (fig. S6). We thus reasoned that the two complexes might share channel activity. To test this conjecture we used an assay previously established in *Xenopus laevis* (*Xenopus*)oocytes (*3*) to assess potential channel activity of the Sr35 resistosome. Co-expression of *Sr35* and *AvrSr35*, but not either alone, generated currents as recorded by two-electrode voltage-clamp (Fig. 3A and B), suggesting that assembly of the Sr35 resistosome is required for the currents. In strong support of this conclusion, two tested *Sr35* mutants, affecting the interface with AvrS35 and abolishing AvrSr35-dependent cell death activity of the receptor *in planta* (Sr35^R730D/R755Q^ and Sr35^W803L/K754G^; see below), lost their ability to produce currents in oocytes (Fig. 3C). Substitutions affecting the acidic inner lining of the funnel formed by α1-helices in ZAR1^E11A^ have been shown to abolish cell death *in planta* and channel activities in oocytes (*2*, *3*). Unexpectedly, both Sr35 channel and cell death activities were tolerant of these analogous acidic residue substitutions (Sr35^E17A/E22A^) (Fig. 3C to F).

**FIG. 3.**
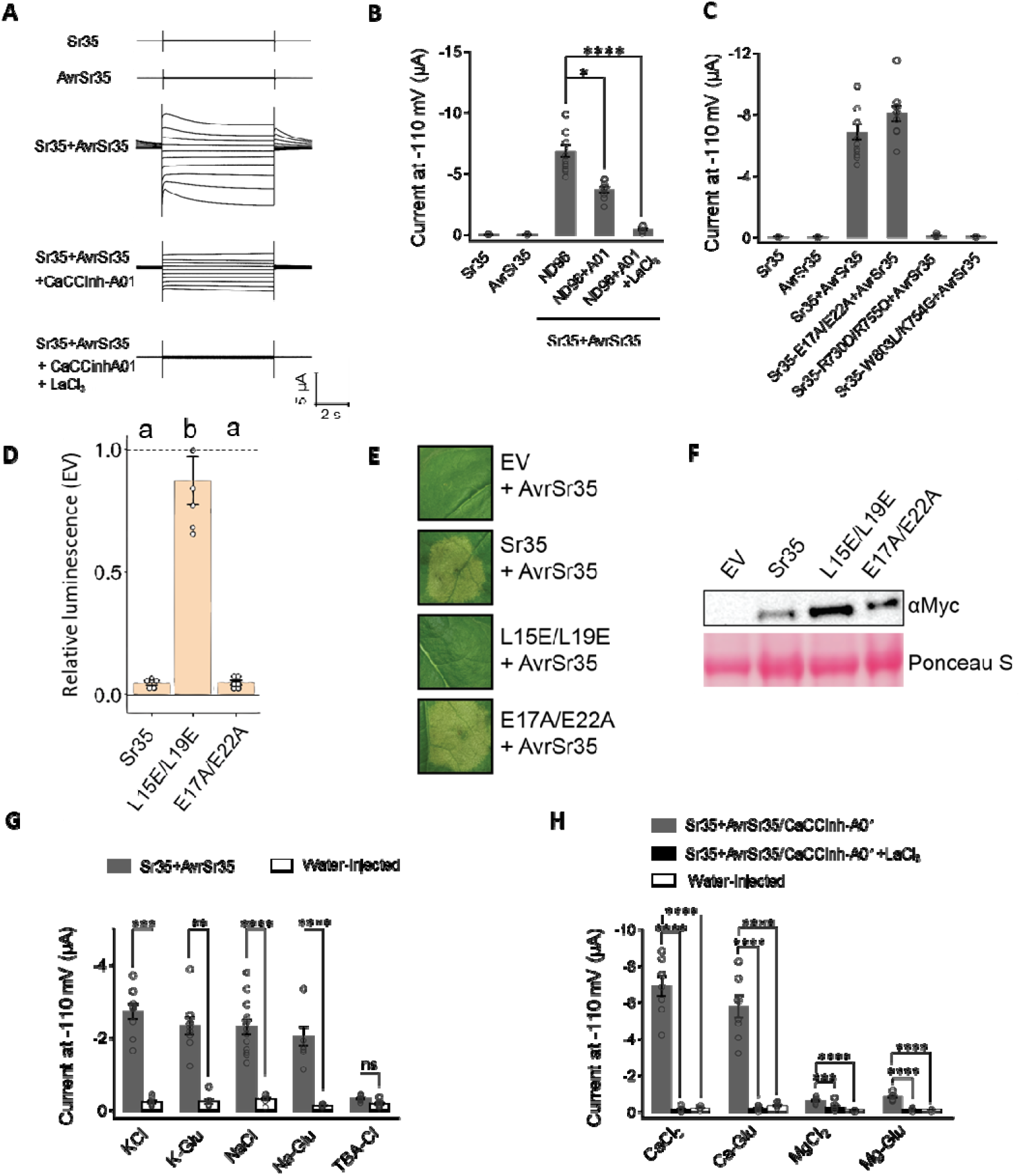
Sr35 resistosome forms a Ca^2+^-permeable non-selective cation channel in *Xenopus* oocytes. (**A** and **B**) TEVC recordings from *Xenopus* oocytes expressing *Sr35, AvrSr35* and *Sr35/AvrSr35*. The effects of CaCCinh-A01 and LaCl_3_ on the *Sr35*-mediated currents were examined. Representative current traces at different voltages from −110 mV to +70 mV in 20 mV increments are shown in (A). Current amplitudes measured at −110 mV are shown in (B). (**C**) Structure-based mutagenesis of Sr35 and AvrSr35 affecting the interface of Sr35 receptor and AvrSr35 effector and Sr35 α1-helix. TEVC recordings were performed in ND96 solution, and current amplitudes measured at −110 mV are shown. (**D**) Wheat protoplast data of Sr35 mutations at α1-helix. Mutagenesis of Sr35 in α1-helix does not abolish cell death in wheat protoplasts when co-expressed with *AvrSr35*. Relative luminescence was used as a readout for cell death. An empty vector treatment was used as the relative luminescence baseline of 1 (mean ± SEM; n = 6). Statistical analysis with one-way ANOVA and Tukey’s test. Treatments labelled with different letters differed significantly (p < 0.05) Additional statistical values reported in methods. (**E**) Tobacco cell death data of Sr35 and Sr35 channel mutants. Representative data is shown from a minimum of eight replicates. (**F**) Western blot of Sr35 and Sr35 channel mutants tested in tobacco. Ponceau S staining was used as a loading control. (**G**) The Sr35 channel is selective for cations. TEVC recordings were performed in various solutions, including KCl (96 mM), K-Gluconate (96 mM), NaCl (96 mM), Na-Gluconate (96 mM), and TBA-Cl (Tetrabutylammonium chloride, 96 mM). (**H**) Cationic currents of CaCl_2_, Ca-Glu, MgCl_2_ and Mg-Glu in presence of CaCCinh-A01 and CaCCinh-A01+LaCl_3_. The divalent cationic currents are blocked by Ca^2+^channel blocker LaCl_3_. (B, G to H) Data are mean ± SEM, n≥8. The asterisks indicate significance of one-way ANOVA analyses and Tukey’s test in (B) and (H), and two-sided Student’s t-tests in (G) *P < 0.033, **P < 0.002, ***P < 0.0002 and ****P < 0.0001.

Xenopus oocytes express endogenous calcium-gated chloride channels (CaCC), and thus, the currents detected in this assay could be confounded by the activity of these native channels. However, addition of the CaCC inhibitor A01 only partially blocked the currents in *Xenopus* oocytes (Fig. 3B) and co-treatment with A01 and the calcium channel blocker LaCl_3_ was required for complete inhibition of electrical activity (Fig. 3B). Together, these results suggest that the Sr35 resistosome has calcium channel activity in *Xenopus* oocytes.

To test whether Sr35 can function as a nonselective cation channel, we tested cation flux in the presence of monovalent solutions of potassium and sodium chloride salts (KCl, NaCl). Similar to ZAR1, co-expression of *Sr35* and *AvrSr35* increased cation flux in oocytes, which was retained for potassium and sodium salts with the immobile gluconate counter-ion (K-Glu, Na-Glu). In contrast, we observed only residual ion flux when a chloride salt of the immobile Tetrabutylammonium was used (TBA-Cl) (Fig. 3G). A comparison of the divalent ions Ca^2+^ and Mg^2+^ (MgCl_2_, CaCl_2_) combined with the *Sr35* and *AvrSr35* co-expression in oocytes, showed that ion flux was significant for calcium but not magnesium (Fig. 3H). This finding supports the conclusion that Sr35 is permeable to calcium. While our collective data strongly suggests that the Sr35 resistosome functions similarly to ZAR1 by forming a nonselective calcium channel, the channel activity of the Sr35 resistosome is tolerant to substitutions of acidic residues predicted to line the inner surface of the channel. Thus, we cannot exclude the possibility that the very N-terminus of the Sr35 resistosome (residues 1 to 21) is structurally and functionally distinct from that of the ZAR1 resistosome.

### The AvrSr35 effector is directly recognised by C-terminal LRRs of Sr35

In the cryo-EM structure, AvrSr35 binds to the very C-terminal ascending lateral side of the LRR domain, supporting a direct recognition mechanism of the effector by Sr35 (Fig. 4A). AvrSr35 is much larger than many other pathogen effectors, but only a small portion of the protein is involved in contact with the Sr35 LRR through charge and shape complementarity (Fig. 4A, fig. S7). Nearly all residues from the ascending lateral side of the last eight LRRs contribute to recognizing AvrSr35, and many of the residues interact with a single helix (α10) of AvrSr35. AvrSr35^Y383^, AvrSr35A^384^, AvrSr35^Y387^ and AvrSr35A^388^ from one α10 side are located at the centre of the Sr35-AvrSr35 interface and make extensive contacts with their respective neighbouring residues in Sr35 (Fig. 4B). Several residues from the loop region C-terminal to α10 form hydrophobic contacts with Sr35^W919^. Similar interactions are also made between AvrSr35^R381^ in the N-terminal side of α10 and Sr35. In addition to the hydrophobic and van der Waals interactions, a large network of hydrogen bonds also mediates the Sr35-AvrSr35 interface, supporting specific recognition of AvrSr35 by Sr35 (individual contacts provided in Fig. 4B).

**FIG. 4.**
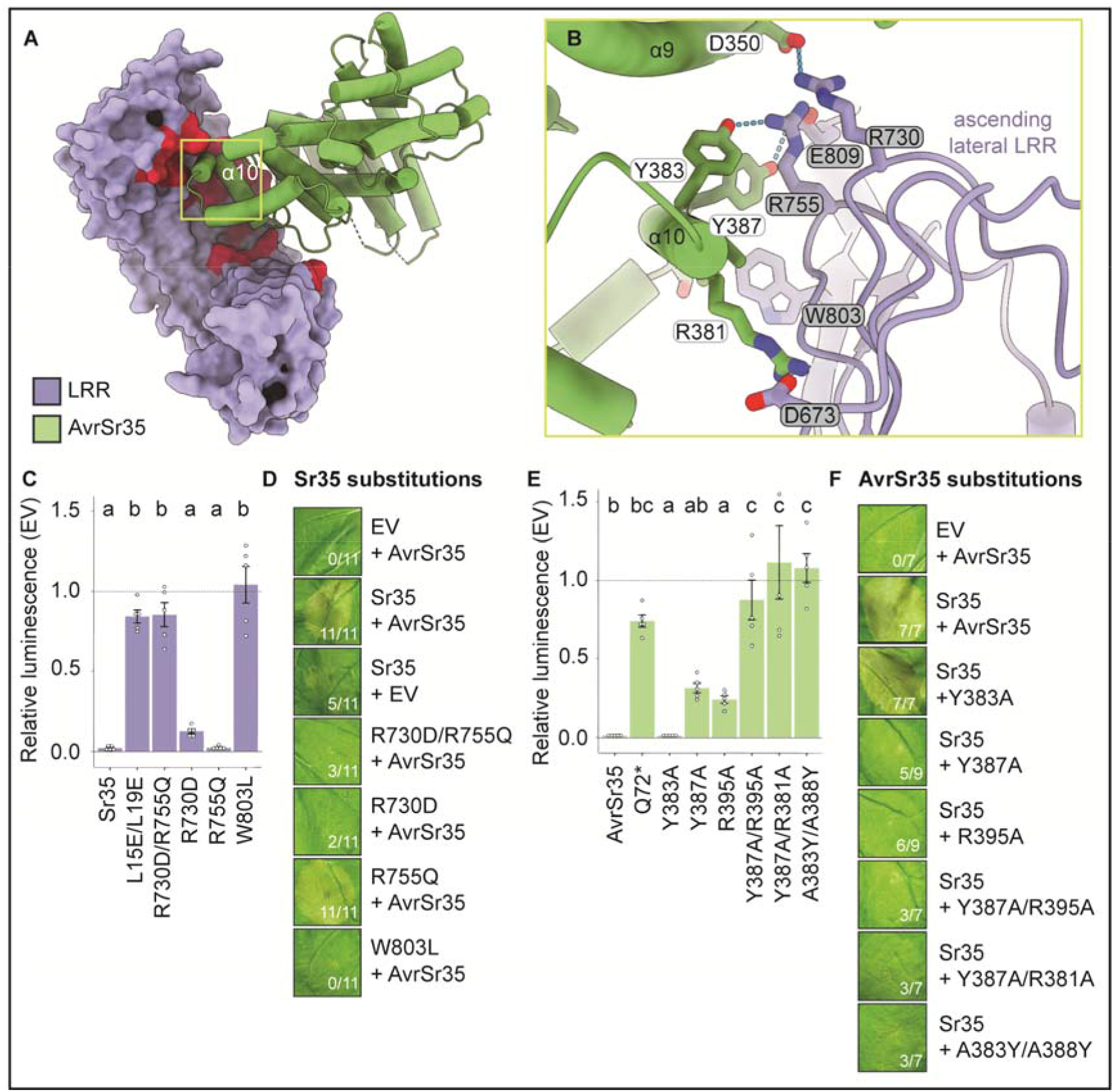
Direct AvrSr35 effector recognition is mediated by Sr35 LRR (**A**) Interface between Sr35 LRR and AvrSr35. Red colour indicates the critical LRR residues that are in a 5 Å radius from AvrSr35. (**B**) Structural detail of Sr35 receptor and AvrSr35 effector interface. Dashed lines represent polar interactions. Grey and white residue label boxes corresponding to Sr35 and AvrSr35 sidechains, respectively. (**C**) Co-transfection of *Sr35* LRR mutants with *AvrS35* in wheat protoplasts. Relative luminescence was used as a readout for cell death. Relative luminescence was used as a readout for cell death. An empty vector treatment was used as the relative luminescence baseline of 1 (mean ±SEM; *n* = 5). Statistical analysis with one-way ANOVA and Tukey’s test. Treatments labelled with different letters differed significantly (*p* < 0.05). Additional statistical values reported in methods. Bar colours correspond to box colours in (B). Bar colours correspond to domain colour code defined in (A). (**D**) *N. benthamiana* cell death data of Sr35 LRR mutations at the receptor-effector interface. Representative data is shown from 11 replicates and scored for leaf cell death. (**E**) Co-transfection of *Sr35* with *AvrS35* mutants in wheat protoplasts. Experimental layout and statistics as in (C). Bar colours correspond to domain colour code defined in (A). (**F**) *N. benthamiana* cell death data of *AvrSr35* mutants co-expressed with *Sr35*. Representative data is shown from 9 replicates and scored for leaf cell death.

To functionally verify the Sr35-AvrSr35 interaction, we first substituted R730, R755, and W803 in Sr35 with their counterparts in the Sr35 homolog of wheat cultivar Chinese Spring(*25*) (here denoted *TaSh1*), that shares 86.5% sequence identity with Sr35 but is derived from a wheat cultivar susceptible to Pgt strains harbouring AvrSr35 (*12*). These W803L or R730D substitutions strongly and weakly suppressed *Sr35*-mediated cell death activity, respectively, when co-expressed with *AvrSr35* in wheat protoplasts (Fig. 4C). In contrast, R755Q had no detectable effect on *Sr35*-induced cell death, but a combination of R730D with R755Q resulted in a full loss of cell death in wheat protoplasts (Fig. 4C). Similar results were obtained when these *Sr35* mutants were assayed in *N. benthamiana* (Fig. 4D and fig. S4C). These data support the Sr35-AvrSr35 interaction as seen in the cryo-EM structure and explain why *Ta*SH1 in susceptible cultivar Chinese Spring is unable to recognize AvrSr35. To further verify specific AvrSr35 recognition by Sr35, we made the following substitutions in the fungal effector at their interface: R383A, Y387A, R395A, Y387A/R395A, Y387A/R381A, A383Y/A388Y, all of which either form hydrogen bonds or salt-bridges with Sr35 LRR (Fig. 4B). The mutations Y387A/R395A, Y387A/R381A and A383Y/A388Y abolished *Sr35*-induced cell death in wheat protoplasts and *N. benthamiana* (Fig. 4E and F, and fig. S4D). In contrast, single mutations of R383A, Y387A, R395A, (Fig. 4E and F, and fig. S4D) and others of AvrSr35 (fig. S8) did not abrogate effector-triggered receptor activation, suggesting that a large portion of the AvrSr35-Sr35 interface is resilient to single mutational perturbations.

### Steric clash between effector and Sr35 NBD mediates receptor activation

To understand how AvrSr35 activates the Sr35 resistosome, we modelled a structure of inactive Sr35 using AlphaFold 2 (*26*). Structural alignment of the LRR domain from the Sr35 resistosome and the LRR from the inactive Sr35 model shows striking similarity (fig. S9A). However, substantial differences exist between the cryo-EM and modelled structures of fulllength Sr35, with structural re-organization within the NOD module (NBD-HD1 relative to WHD) (fig. S9B). This is reminiscent of the activation of the ZAR1 resistosome, which involves an allosteric mechanism (*1*, *2*). Modelling of AvrSr35 onto the LRR domain of the predicted structure of inactive Sr35 shows substantial overlap between the effector and Sr35 NBD (fig. S10). This ‘steric clash’ suggests that AvrSr35 binding dislodges the NBD, allowing subsequent nucleotide exchange for further ATP-triggered allosteric changes in the receptor and assembly of the Sr35 resistosome. In this context, effector binding to the ascending lateral side of the LRR domain appears a conserved mechanism of plant NLRs (fig. S11). Together, these results support a conserved allosteric mechanism underlying activation of the Sr35 and ZAR1 resistosomes.

### Structure-enabled gain-of-function engineering of orphan CNLs

To test whether these insights into the evolutionary conservation of CNL resistosomes can be harnessed for the design of novel receptors with altered function, we first chose two closely related *Sr35* homologs (*Sh*) of unknown resistance function in bread wheat (*Triticum aestivum; TaSh1*) and in the sister species barley (*<u>H</u>ordeum <u>v</u>ulgare; HvSh1*). We generated hybrid receptors of *TaSh1* and *HvSh1* in which the LRR domain, including the highly conserved WHD α4-helix, was substituted by the equivalent fragment of Sr35 (termed *TaSh1^Sr35LRR^* and *HvSh^Sr35LRR^;* Fig. 5A and fig. S12). Unlike wild-type *TaSh1* or *HvSh1* genes both hybrid receptors mediated *AvrSr35*-dependent cell death in wheat leaf protoplasts prepared from cultivar Chinese Spring and when expressed in leaves of *N. benthamiana* (Fig. 5B to D), indicating neofunctionalization of the orphan receptors.

**FIG. 5.**
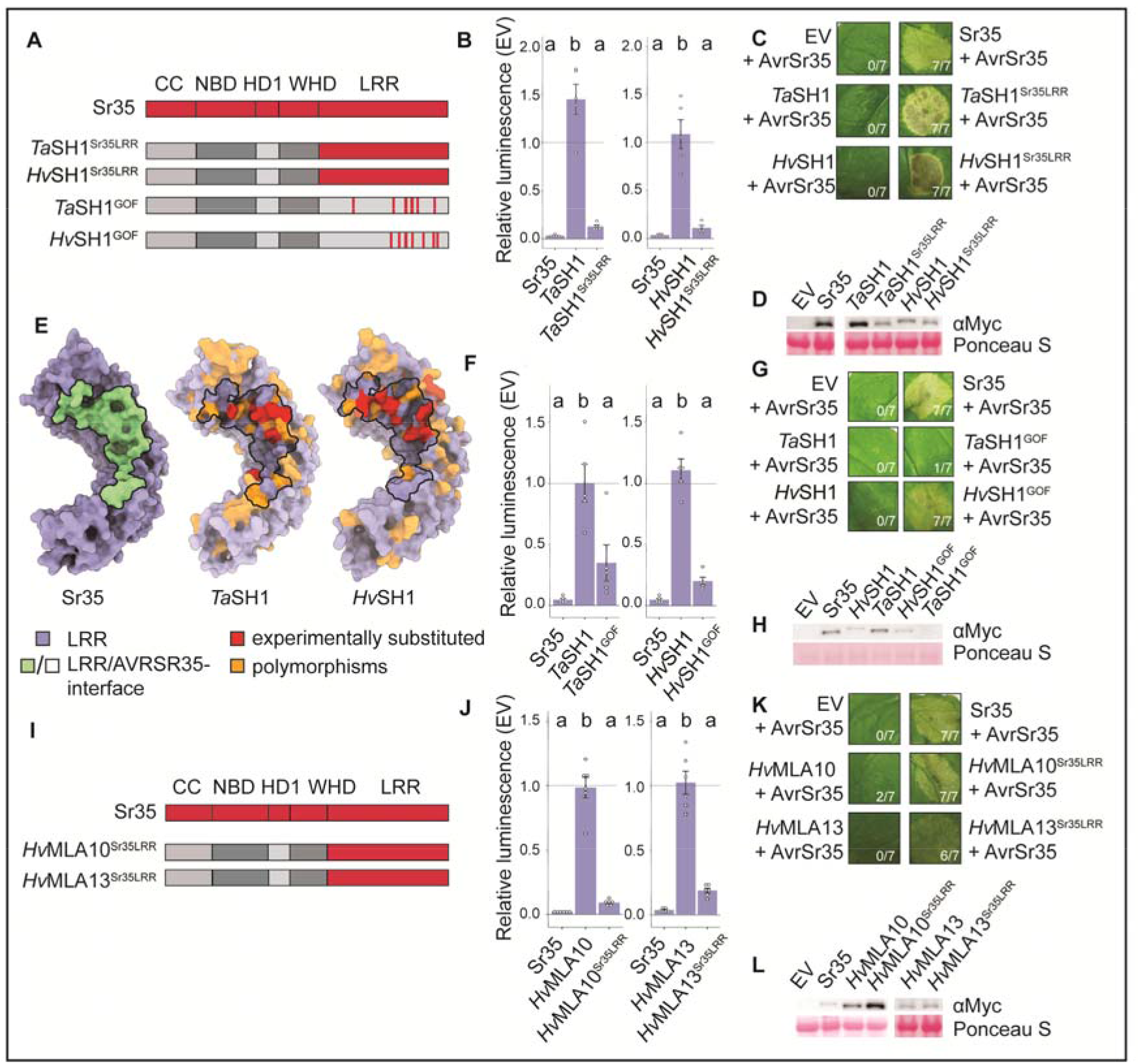
Structure-guided neofunctionalization for AvrSr35-recognition of orphan CNLs and CNL receptor hybrids. (**A**) Graphic illustration of Sr35 domain structure and hybrid receptors made on the basis of *Sr35 homologs* (*Sh*)of unknown function in bread wheat (*Triticum aestivum; TaSh1*) and in the sister species barley (*Hordeum vulgare; HvSh1*). Sr35 LRR domain (red) was used to substitute *Ta*SH1 and *Hv*SH1 LRR domain including α4-helix of the conserved WHD domain (*Ta*SH1^Sr35LRR^ and *HV*SH1^Sr35LRR^). Gain-of-function (GOF) receptor variants (*Ta*SH1^GOF^ and *Hv*SH1^GOF^ were created based on sequence polymorphisms between Sr35, *Ta*SH1 and *Hv*SH1. (**B**) Wheat protoplast transfections of *Ta*SH1^Sr35LRR^ and *Hv*SH1^Sr35LRR^ and controls. Relative luminescence was used as a readout for cell death. An empty vector treatment was used as the relative luminescence baseline of 1 (mean ±SEM; *n* = 5). Statistical analysis with one-way ANOVA and Tukey’s test. Treatments labelled with different letters differed significantly (*p* < 0.05). Additional statistical values reported in methods. (**C**) Tobacco cell death data of *Ta*SH1 and *Hv*SH1 expression. Representative data is shown from seven replicates and scored for leaf cell death. (**D**) Western blot of chimeras tested in *N. benthamiana*. Pooled samples from three technical replicates. Ponceau S staining as a loading control. Left and right part of one blot were merged into a composite image. (**E**) Cryo-EM structure of Sr35 and predicted structures of *Ta*SH1 and *Hv*SH1 LRR domains (Jumper et al. 2021). All polymorphisms of *Ta*SH1 and *Hv*SH1 relative to Sr35 are shown (orange). Structural alignment was used to identify polymorphisms at LRR-AvrSr35 interface (red) and mutated. (**F**) Wheat protoplast transfections of *Ta*SH1^GOF^ and *Hv*SH1^GOF^ and controls. Relative luminescence was used as a readout for cell death. Experimental design and statistical analysis as in (b) (*n* = 5). (**G**) Tobacco cell death data of *Ta*SH1^GOF^ and *Hv*SH1^GOF^ expression. Representative data is shown from seven replicates and scored for leaf cell death. (**H**) Western blot of GOF experiment tested in *N. benthamiana*. Pooled samples from three technical replicates. Ponceau S staining as a loading control. (**I**) Graphical illustration of MLA proteins used for chimer design. Sr35 LRR domain (red) was used to substitute *Hv*MLA10 and *Hv*MLA13 LRR including α4 of the conserved WHD domain (*HvMla10^Sr35LRR^* and *HvMLA13^Sr35LRR^*). (**J**) Wheat protoplast transfections of *HvMla10^Sr35LRR^* and *HvMLA13^Sr35LRR^* compared to *Mla* WT and *Sr35* controls. Experimental layout and statistical analysis as in (B). (*n*=6) (**K**) Tobacco cell death data of *HvMla10^Sr35LRR^* and *HvMla13^Sr35LRR^* constructs. Representative data is shown from seven replicates and scored for leaf cell death. (**L**) Western blot of MLA hybrid receptor experiment tested in *N. benthamiana*. Pooled samples from 3 technical replicates. Ponceau S staining as a loading control. Left and right part of two independent blots were merged into a composite image.

Due to the high sequence similarity of *Ta*SH1 and *Hv*SH1 with *T. monococcum* Sr35 (86.5% and 86.4% amino acid sequence identity to Sr35, respectively) we reasoned that single amino acid substitutions in the LRR domains of the homologs might be sufficient to enable detection of AvrSr35. Combined structural model and protein sequence alignments indicated that the AvrSr35-interacting residues of Sr35 are polymorphic in *Ta*SH1 and *Hv*SH1 (Fig. 5E and fig. S13). The alignments identified several residues in the LRR domains of *Ta*SH1 and *Hv*SH1 that likely hinder effector binding at the modelled interface. Accordingly, we generated *TaSH1* and *HvSH1* variants encoding receptors with eight or ten substitutions in the LRR, respectively (*Ta*SH1^D609G/Y728F/D731R/I754K/Q755R/L804W/Q810E/R857W^ and *Hv*SH1^Y727F/Q801E/G754K/Q752P/Q755R/R809E/W835I/R856W/917D/P919W^;designated for simplicity *Ta*SH1^GOF^ and *Hv*SH1^GOF^, Fig. 5A). Unlike wild-type *HvSh1*, *HvSh1^GOF^* mediated a clear cell death response in wheat protoplasts and *N. benthamiana* when co-expressed with the effector *AvrSr35* (Fig. 5F to H). *Ta*SH1^GOF^ induced a striking cell death phenotype in wheat protoplasts, but not *N. benthamiana* (Fig. 5F to H), which is likely due to undetectable *Ta*SH1^GOF^ protein in the heterologous *N. benthamiana* expression system (Fig. 5H). The findings suggest that single amino acid substitutions mimicking the effector binding region of Sr35 are sufficient for neofunctionalization of these orphan receptors. The relatively small number of nucleotide changes needed in order to enable *Ta*SH1 to detect AvrSr35 makes it feasible to introduce such changes in elite bread wheat by gene editing. In this way, generating varieties that are resistant to *Pgt* Ug99 (*27*–*31*) provides an alternative strategy to transgene-mediated *Sr35* transfer from *T. monococcum* to bread wheat (*14*, *32*).

Next, we investigated whether the Sr35 LRR domain, transferred to more distant CNLs (~45% amino acid sequence identity) in the sister species barley, can generate functional hybrid receptors. We chose barley *Hv*MLA10 and *Hv*MLA13, known to confer isolate-specific immunity against the barley powdery mildew fungus (*33*), *Blumeria graminis* f sp *hordei* (*Bgh*), as templates to engineer AvrSr35 recognition (Fig. 5I). The ascomycete *Bgh* effectors recognized by *Hv*MLA10 and *Hv*MLA13 lack sequence similarity to AvrSr35 from the basidiomycete *Pgt*. Using the above reasoning for hybrid receptor generation of *Sr* homologs, the LRR domains of *Hv*MLA10 and *Hv*MLA13 were replaced by the Sr35 LRR. The two resulting hybrid receptors, *HvMla10 ^Sr35LRR^* and *HvMla13 ^Sr35LRR^*, induced cell death when co-expressed with *Pgt AvrSr35* in wheat protoplasts and *N. benthamiana* (Fig. 5J to L). This finding supports our hypothesis that a combination of effector binding to the LRR and steric clash of the effector with the NBD is needed for CNL activation, as exemplified here for hybrid receptors where the AvrSr35 effector is predicted to clash with the MLA NBD.

## DISCUSSION

Our results, together with earlier data, strongly suggest that CNLs are evolutionarily conserved among monocotyledonous and dicotyledonous plants. Two lines of evidence support this idea: our structural elucidation of the wheat Sr35 resistosome and its similarity to the previously reported Arabidopsis ZAR1 resistosome structure (*2*), and the functional interspecies hybrid CNL receptors generated in this study from wheat Sr35 and barley. This conservation extends to the non-selective cation flux across membranes enabled by pentamerization, although it is possible that ion selectivity and channel dynamics differ between individual CNLs, including the channel activity of so-called helper NLRs acting downstream of canonical plant NLRs (*34*). Reconstitution of effector-dependent Sr35-triggered cell death in insect cells indicates that regulated channel activity is sufficient to recapitulate plant CNL-mediated cell death in eukaryotic cells of another kingdom. It is possible that plant CNL pore formation and ion flux trigger and intersect with intrinsic cell death pathways in animals, e.g. Apaf-1 apoptosome-mediated developmental and stress-induced cell death (*35*, *36*). Although the components needed for cell death downstream of CNL channel activity in plants remain to be identified, the evolutionary conservation of channel activity rationalizes how highly diverse pathogen signals activate a shared set of downstream responses. This is reminiscent of the highly conserved NADase activity encoded by the TIR domain, which converts the presence or activity of diverse pathogen molecules into TNL-triggered immune signals (*37*, *38*).

Our study also uncovers the mechanism by which direct or indirect recognition of pathogen effectors results in the formation of the conserved pentameric scaffold encoding channel activity. Indirect recognition of a bacterial pathogen effector by ZAR1 results in a conformational change of the NBD, allosterically promoting exchange of ADP with ATP/dATP for full receptor activation. Our data support a similar mechanism for Sr35 activation by direct recognition of AvrSr35. These results lend further support to the notion that exchange of ADP with ATP/dATP is widely involved in the activation of NLRs. Although AvrSr35 is absolutely required to initiate Sr35 activation, the effector makes no contribution to oligomerization of the Sr35 resistosome, which is principally mediated by the conserved NBD. This is also true for the assembly of the ZAR1 resistosome and the Apaf-1 apoptosome in animals (*2*, *39*). It seems that recognition of diverse pathogen effectors by the polymorphic LRRs release the conserved NBD to mediate NLR oligomerization.

A third plant NLR recognition mechanism involves a combination of direct and indirect recognition through the incorporation of effector target domains (integrated decoys, ID) into the NLR domain architecture, designated *NLR-IDs*, representing approximately 10% of all *NLR* genes of a plant species (*40*). Crystal structures of the ID in complex with the bound pathogen effector have been resolved for two NLR-IDs, enabling structure-informed ID engineering to extend pathogen strain-specific NLR recognition (*41*–*43*). How the corresponding full-length NLR-ID receptors are activated, including a potential steric clash with their NBD is unclear due to a lack of full-length receptor structures. This is further complicated as NLR-IDs, which directly recognize effectors, typically function with canonical NLRs as interacting pairs (*40*). Direct recognition of pathogen effectors by plant NLRs can be rapidly circumvented by polymorphisms of effector residues at the effector–NLR interface, particularly as a pathogen and its host plant evolve at different time scales. Virulent isolates of *Pgt* within and beyond the Ug99 lineage have escaped the recognition of at least one of the recently cloned *Sr* genes, including single amino acid changes in the effector (*44*). For example, a *Pgt* isolate with combined virulence against *Sr35* and *Sr50* caused an epidemic in Sicily in 2017 (*45*). Our findings allow the prediction not only of AvrSr35 substitutions that might escape Sr35 recognition, but also substitutions in the Sr35 LRR that can physically ‘recapture’ such effector variants. More generally, the evolutionarily conserved plant CNL resistosome architecture with its conserved function highlights the future potential of structure-guided NLR engineering for crop improvement.

## MATERIALS AND METHODS

### Protein expression, purification and negative staining

Codon optimized *Sr35^L15E/L19E^* and *AvrSr35* genes were cloned into the *pFastBac1* vector (Invitrogen) with an N-terminal 6×His-SUMO tag and an N-terminal GST tag, respectively. The constructs were transformed into EMBacY (*46*) competent cells for recombinant bacmid DNA generation. Recombinant baculovirus was generated by initial lipofection with Xtreme gene reagent (Roche) of Sf21 insect cells (Invitrogen). Baculovirus was generally amplified to the P2 generation before protein expression. *Sr35^L15E/L19E^* and *AvrSr35* were co-expressed in Sf21 insect cells, 50 mL of each virus was used per 1 L of culture. After expression of protein at 28 °C for 48 h, the insect cells were harvested and resuspended with buffer containing 50 mM Tris pH 8.0, 150 mM NaCl, 0.05% Triton X-100 and 5% glycerol. The cell lysates generated by sonication were centrifuged at 13,000 *rpm* for 1.5 h, then the supernatant was collected. The protein complex was purified with Glutathione Sepharose 4B (GS4B) resin. After binding to the Glutathione agarose twice, the agarose was washed with three column volumes of resuspension buffer, and the tagged protein complex was treated with GST-tagged PreScission protease at 4°C overnight to remove GST and 6xHis-SUMO tags simultaneously and to elute Sr35-AvrSr35 protein complex. The digested protein complex was subjected to HiLoad superpose 6 column (GE) in buffer containing 50 mM Tris pH 8.0, 100 mM NaCl and 0.01% triton X-100. Pooled peak fractions were used for cryo-EM sample preparation.

### Cryo-EM sample preparation and data collection

The Sr35-AvrSr35 complex grids were prepared for cryo-EM analysis. Holy carbon grids (Quantifoil Au 1.2/1.3, 300 mesh) were glow-discharged for 30 s at medium level in HarrickPlasma after 2 min evacuation. The purified Sr35-AvrSr35 protein was concentrated to ~0.5 mg/mL and 3 μL of sample were applied to the grid. The grids were blotted for 2–3 s using a pair of filter papers (55 mm, Ted Pella Inc.) at 8°C with 100% humidity and flash-frozen in liquid ethane using an FEI Vitrobot Marked IV. Stacks of Sr35-AvrSr35 cryo-EM samples were collected by a Titan Krios microscope operated at 300 kV, equipped with a K3 Summit direct electron detection camera (Gatan) using EPU 2 (Thermo Fisher Scientific) at Zhengzhou University. Micrographs were recorded at 81,000× magnification corresponding to 1.1 Å/pixel. The defocus ranged from −1.5 μm to −2.0 μm. Each image stack contains 32 frames recorded every 0.11 s for an accumulated dose of ~50e/Å^2^ and a total exposure time of 3.5 s. A second data set from an independent protein purification was recorded at EMBL Heidelberg with the following parameters: Titan Krios microscope operated at 300 kV, equipped with a K3 Summit direct electron detection camera (Gatan), 50e/ Å^2^, 40 frames/stack.

### Image processing and 3D reconstruction

All micrographs of the Sr35-AvrSr35 complex were 2 × 2 binned, generating a pixel size of 1.1 Å. MotionCor2 program was used to perform Motion correction (*47*). Contrast transfer function (CTF) parameters were estimated by CTFFIND4 (*48*). Based on the CTF estimations, 5,292 micrographs were manually picked and were further processed in RELION3.1 (*49*).

1,608,441 particles were picked using Laplacian-of-Gaussian auto picking and then subjected to several rounds of 2D classification (*50*, *51*). Every round of 2D classification performed 25 iterations with regularisation parameter T=2 and number of classes=100 to remove bad particles. The particles with the best quality were used to generate the initial model using *ab initio* calculation from RELION3.1. Then 698,386 particles were imported into 3D classification with C1 symmetry. There were five Sr35 molecules in the complex, each of them was bound to a ligand (AvrSr35). C5 symmetry was used in the following 3D refinement. After CTF refinement and post-processing, the resolution of the Sr35-AvrSr35 complex reconstruction was 3.0 Å. The resolution was estimated by the gold-standard Fourier shell correlation (FSC) = 0.143 criterion (*52*). Local resolution distribution was evaluated using RELION (*53*).

In the reconstruction above, the LRR and AvrSr35 portions were more flexible than the other parts of the Sr35-AvrSr35 complex. To improve the density of the more flexible portions, a procedure reported before by Bai *et al*. 2015 was employed (*54*). The final refined particles were expanded with C5 symmetry. A local mask was generated using USCF Chimera (*55*). Expanded particles and local mask were subjected to 3D classification without alignment. Finally, 476,069 particles were used for 3D auto-refinement and CTF refinement. A final resolution of 3.33□Å was achieved after postprocessing. For our second dataset, one third of the micrographs were analysed the same way and resulted in the same overall structure at a resolution of 3.4 Å. The resulting model was not used further for model building.

### Model building and refinement

The final density map was obtained by merging the global map and the local map which contained LRR and AvrSr35, using ‘combine_focused_map’ in PHENIX (*56*). The model of the Sr35-AvrSr35 complex was manually built in COOT(*57*) based on the global and the local maps. The generated model was refined against the combined Sr35-AvrSr35 EM density using real space refinement in PHENIX with secondary structure and geometry restraints (*57*). Model statistics can be found in Extended Data Table 1.

### Transient gene expression assays in wheat protoplasts

Seedlings of the wheat cv. Chinese Spring were grown at 19°C, 70% humidity and under a 16 h photoperiod. Protoplasts were isolated from the leaves and transfected as described by Saur *et al*. 2019 (*22*). The coding sequence of *TaSh1* (NCBI XP_044359492.1) and *HvSh1* (NCBI KAE8803279.1) were generated by gene synthesis based on wild-type codons (GeneArt, Invitrogen). The coding sequence of all tested receptor constructs, or an empty vector (EV) as negative control, were expressed from *pIPKB002* vector (*58*) containing the strong ubiquitin promoter. Receptors were co-expressed with *AvrSr35* in *pIPKB002*. In addition, co-transfection of *pZmUBQ:LUC*(*59*) facilitated the expression of the *LUC* reporter construct. Each treatment was transfected with 4.5 μg of *pZmUBQ:LUC* and 5 μg of *pIPKb002:AvrSr35*. Quantities of receptor-encoding *pIPKb002* plasmid were varied for each construct in an effort to minimize cell death due to (receptor) toxicity-mediated cell death (*EV* 8 μg; *Sr35* and *Sr35 mutants* 2 μg; *AvrSr35* and *AvrSr35 mutants* 5 ug; *HvMla10*, *HvMla13*, *HvMla10^Sr35LRR^*, *HvMla13^Sr35LRR^*, *TaSh1*, *TaSh1*, *TaSh1^GOF^*, *TaSh1^GOF^* 8 μg; *TaSh1^Sr35LRR^* and *TaSh1^Sr35LRR^* 2 μg). A maximum of two technical replicates were completed with the same batch of wheat seedlings. Luminescence was measured using a luminometer (Centro, LB960). Relative luminescence was calculated by dividing the absolute luminescence value by that of the corresponding *EV* treatment (EV = 1).

### Transient expression and western blotting in tobacco

For *N. benthamiana* transient expression, *Sr35*, *Sr35* mutants, *AvrSr35* and *AvrSr35* mutants were cloned into the *pDONR* vector (Invitrogen). The obtained plasmids of *Sr35* and *Sr35* mutants were recombined by an LR reaction into *pGWB517-4×Myc* with a C-terminally fused 4×Myc epitope tag (*60*), while *AvrSr35* and *AvrSr35* mutants were recombined into the *pXCSG-mYFP*(*61*) vector with a C-terminally fused mYFP epitope tag. After being verified by Sanger sequencing, all the constructs were transformed into *Agrobacterium tumefaciens* GV3101 pMP90RK by electroporation. Transformants were grown on LB media selection plates containing rifampicin (15 mg/mL), gentamycin (25 mg/mL), kanamycin (50 mg/mL), and spectinomycin (50 mg/mL) for transformants harbouring *pGWB517-4×Myc* or carbenicillin (50 mg/mL) for *pXCSG-mYFP*.

Individual *Agrobacterium* transformants were picked and cultured in LB medium containing respective antibiotics in the abovementioned concentration. After shaking culture at 28°C for 16 h, the culture was harvested at 3,800 rpm for 10 min and resuspended with infiltration buffer containing 10 mM MES pH 5.6, 10 mM MgCl_2_ and 150 μM acetosyringone. The OD_600_ of *AvrSr35* and *AvrSr35* mutant strains was adjusted to 1.0. For *Sr35* and *Sr35* substitution mutants, the OD_600_ was adjusted to 0.15. Hybrid receptor bacterial strains (*HvMla10^Sr35LRR^, HvMla13^Sr35LRR^, TaSh^Sr35LRR^, HvSh^Sr35LRR^*) were adjusted to an OD_600_ of 0.6. In the hybrid receptor gain-of-function experiment, OD_600_ of *TaSh1, HvSh1, TaSh1^GOF^* and *HvSh1^GOF^* bacterial strains was adjusted to 1.8 without resulting cell death in co-expression of *TaSh1* and *HvSh1* when co-expressed with *AvrSr35*. After dilution, all the cell suspensions were incubated at 28°C for 1 h at 200 rpm. Construct expression was conducted in leaves of four-week-old *N. benthamiana* plants via *Agrobacterium-mediated* transient expression assays. For phenotypic experiments, *Agrobacteria* cultures expressing receptor constructs, or the respective receptor mutants, were co-infiltrated with *AvrSr35*, or its mutants, at 1:1 ratio using a syringe. As a control, either receptor or effector bacterial strains were replaced with *Agrobacteria* transformed with EVs. Phenotypic data was recorded at day 3 after infiltration.

*Agrobacterium-mediated* transient expression assays for protein detection were conducted as described above. The infiltrated leaves were harvested at 24 h after infiltration, flash-frozen in liquid nitrogen and ground to powder using a Retsch grinder. Plant powder was mixed with 4xLämmli buffer in 1:2 ratio. Five microliters were loaded onto 10% SDS PAGE. After transfer to PVDF membrane, protein was detected using monoclonal mouse Anti-myc (1:7,500; R950-25, Thermofisher), polyclonal rabbit anti-GFP (1:7,500; ab6556, Abcam), polyclonal goat anti-mouse IgG-HRP (1:5,000; Santa Cruz Biotechnology, sc2005) and polyclonal donkey anti-rabbit IgG-HRP (1:5,000; Santa Cruz Biotechnology, sc2313) antibodies. Protein was detected using SuperSignal West Femto:SuperSignal substrates (ThermoFisher Scientific) in a 1:1 ratio.

### Electrophysiology

The two-electrode voltage clamp (TEVC) recordings were conducted as previously described (*3*). The cDNAs of *Sr35*, or *Sr35* mutants, and *AvrSr35* were cloned into the *pGHME2* plasmid for expression in *Xenopus* oocytes. cRNAs for all constructs were transcribed using T7 polymerase. Ovarian lobes were obtained from adult *Xenopus laevis* under anaesthesia. Both the amount of cRNA injected and the oocyte incubation time were optimized to minimize toxicity caused by the assembled Sr35 resistosome. Isolated oocytes were coinjected with 0.5 ng cRNA of *Sr35* (WT and mutants) and *AvrSr35*. Oocytes were then incubated at 18°C for approximately 4 h in ND96 buffer (96 mM NaCl, 2.5 mM KCl, 1 mM MgCl_2_, 1.8 mM CaCl_2_, 5 mM HEPES pH 7.6) supplemented with 10 μg l^-1^ penicillin and 10 μg l^-1^ streptomycin. TEVC measurements were performed between 4–7 h later after injection. Water-injected oocytes served as controls.

Two-electrode voltage-clamp recordings were performed using an OC-725C oocyte clamp amplifier (Warner Instruments) and a Digidata 1550 B low-noise data acquisition system with pClamp software (Molecular Devices). The microelectrode solutions contained 3 M KCl (electrical resistance of 0.5-1 MΩ), and the bath electrode was a 3 M KCl agar bridge. To eliminate the chloride currents mediated by endogenous Ca^2+^-activated chloride channels in Xenopus oocytes, the ND96 recording solution was supplemented with 200 μM CaCC inhibitor (CaCCinh)-A01, and the oocytes were pre-incubated 5–10 min before measurement. To test the channel blocking effect of LaCl_3_, the oocytes were pre-incubated for 5–10 min in the recording solutions supplemented with 200 μM CaCCinh-A01 and 100 μM LaCl_3_ before measurement. For the recordings in Fig. 3G, the various recording solutions were as followings: KCl (96 mM), K-Gluconate (96 mM), NaCl (96 mM), Na-Gluconate (96 mM), and TBA-Cl (Tetrabutylammonium chloride, 96 mM). All solutions contained 5 mM HEPES pH 7.6, and 1 mM MgCl_2_ or Mg-Gluconate. For the recordings in Fig. 3H, the various recording solutions were as follows: CaCl_2_ (12 mM), Ca-Gluconate (12 mM), MgCl_2_ (12 mM), and Mg-Gluconate (12 mM). All solutions contain 5 mM HEPES pH 7.6, and 1 mM MgCl_2_ or Mg-Gluconate. The treatments of CaCCinh-A01 and LaCl_3_ were conducted as above. Voltage-clamp currents were measured in response to voltage steps lasting 7.5 s and to test potentials ranging from −110 mV to +70 mV, in 20 mV increments. Before each voltage step, the membrane was held at 0 mV for 1.60 s, and following each voltage step, the membrane was returned to 0 mV for 2 s. I–V relations for Sr35 resistosome channels were generated from currents that were measured 0.2 s before the end of each test voltage step. Three independent batches of oocytes were investigated and showed consistent findings. Data from one representative oocyte batch are shown.

### Statistical analysis

Data distribution for each protoplast transfection experiment was subjected to the Shapiro-Wilk normality test. All experiments were found to be normally distributed. An analysis of variance (ANOVA) and subsequent Tukey’s honest significance difference (HSD) test was completed for each experiment. Treatments found to be significantly different were labelled with different letters (α = 0.05). F-values and associated p-values for each figure are as follows: Fig. 2G: (F(6,28)=[7.778], p = 5.52e-05); Fig. 2I: (F(5,24)=[95.05], p = 5.16e-15); Fig. 3D: (F(3,20)=[64.52], p = 1.84e-10); Fig. 4C: (F(5,24)=[65.69], p = 3.27e-13); Fig. 4E: (F(7,32)=[20.54], p = 3.62e-10); Fig. 5B TaSH1: (F(2, 12)=[75.58], p = 1.58e-07); Fig. 5B HvSH1: (F(2, 12)=[43.74], p = 3.08e-06); Fig. 5F TaSH1: (F(2, 12)=[15.68], p = 0.00045); Fig. 5F HvSH1: (F(2, 12)=[101.9], p = 2.95e-08); Fig. 5J_HvMLA10: (F(2, 15)=[131.7], p = 3.05e-10); Fig. 5J HvMLA13: (F(2, 15)=[105.2], p = 1.5e-09).

## Ethics declarations

### The authors declare no competing interests

*Xenopus laevis* experiments were conducted at State Key Laboratory of Molecular Developmental Biology, Institute of Genetics and Developmental Biology, Chinese Academy of Sciences, Beijing, China; Innovative Academy of Seed Design, Chinese Academy of Sciences, Beijing 100101, China, and in accordance with local legislation.

## Acknowledgments

Zhengzhou University and EMBL Heidelberg are acknowledged for their assistance with Cryo-EM data acquisition. We thank Ulla Neumann (MPIPZ) and Felix Babatz (CECAD) for TEM support, and Neysan Donnelly (MPIPZ) for manuscript polishing.

## Funding

The Alexander von Humboldt Foundation (a Humboldt professorship to J.C.), the Max-Planck-Gesellschaft (P.S.-L. and a Max Planck fellowship to J.C.), Deutsche Forschungsgemeinschaft SFB-1403-414786233 (J.C. and P.S.-L.) and Germany’s Excellence Strategy CEPLAS (EXC-2048/1, Project 390686111) (J.C. and P.S.-L.). iNEXT-Discovery for funding Cryo-EM data set collection at EMBL Heidelberg (PID 16414 to A.F. and J.C.), the National Key Research and Development Program of China 2021YFA1300701 (Z.H.), the Strategic Priority Research Program of the Chinese Academy of Sciences (XDA24020305 to Y.-H.C.) and the National Key Research and Development Program of China (2020YFA0509903 to Y.-H.C.)

## Notes

### Competing Interest Statement

The authors have declared no competing interest.

