## Supplementary Materials for "A wheat resistosome defines common principles of immune receptor channels"

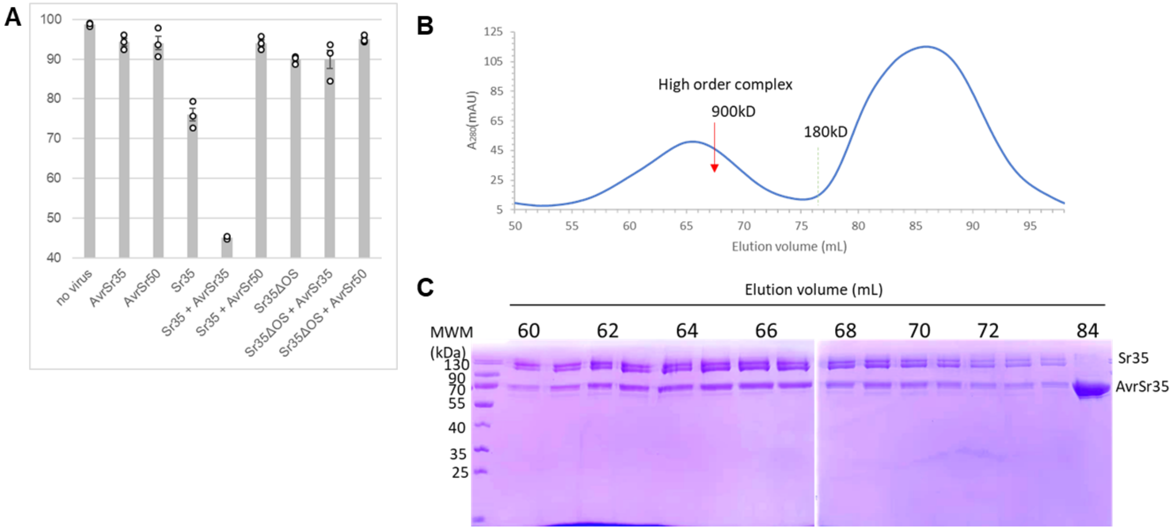


**Fig. S1** Sr35-AvrSr35 expression in Sf21 insect cells. (**A**) Cell viability data in Sf21 insect cells. *Sr35* constructs carrying amino-terminal 6xHis-Sumo-tag and *Avr35* constructs carrying C-terminal GST-tag. Cell viability was measured using trypan blue stain. N=3 biological replicates, error bars=standard error. (**B** and **C**) Purification of Sr35-AvrSr35 resistosome. Purification of Sr35-AvrSr35 resistosome by SEC (B) and SDS PAGE analysis on SEC fractions (C). Numbers represent elution volumes in mL.

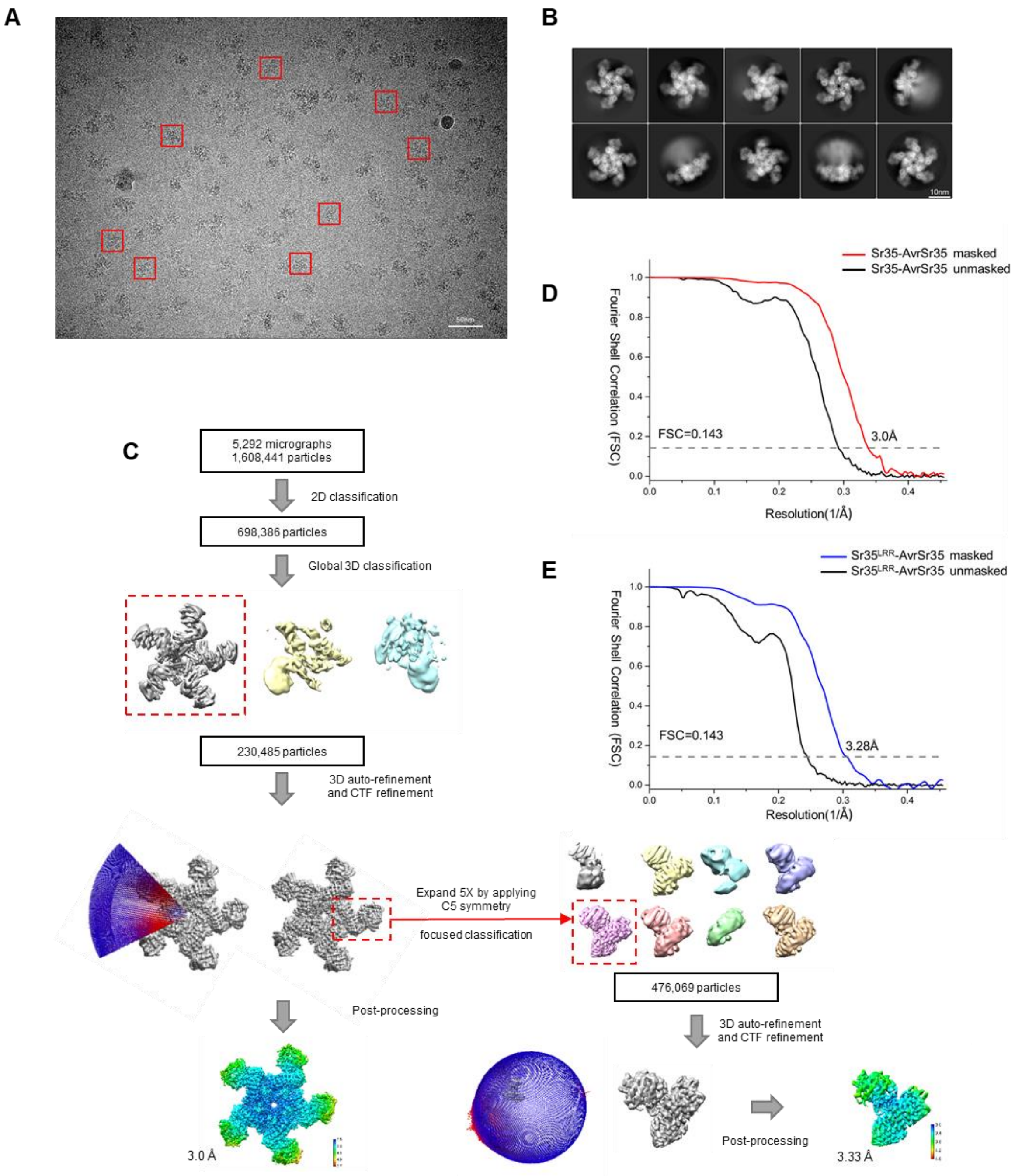


**Fig. S2** Flowchart of Sr35-AvrSr35 3D reconstruction. (**A**) Representative cyro-EM micrograph of Sr35-AvrSr35 complex. (**B**) Representative 2D class averages of Sr35-AvrSr35 complex. (**C**) Flowchart of cryo-EM data processing and Sr35-AvrSr35 3D reconstruction. (**D**) FSC curves at 0.143 of the final model of Sr35-AvrSr35 complex. (**E**) FSC curves at 0.143 of the final model of Sr35LRR-AvrSr35.

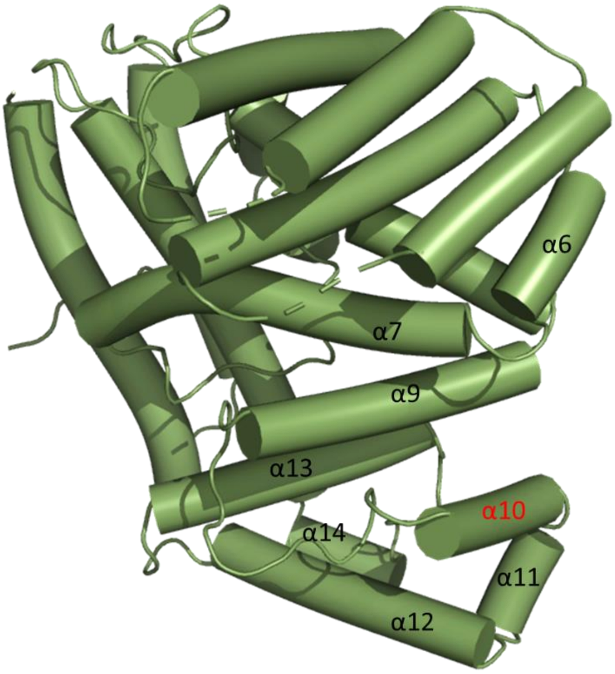


**Fig. S3** AvrSr35 structure from the Sr35 resistosome. α10-helix (red) is involved in most extensive contacts with Sr35 LRR.

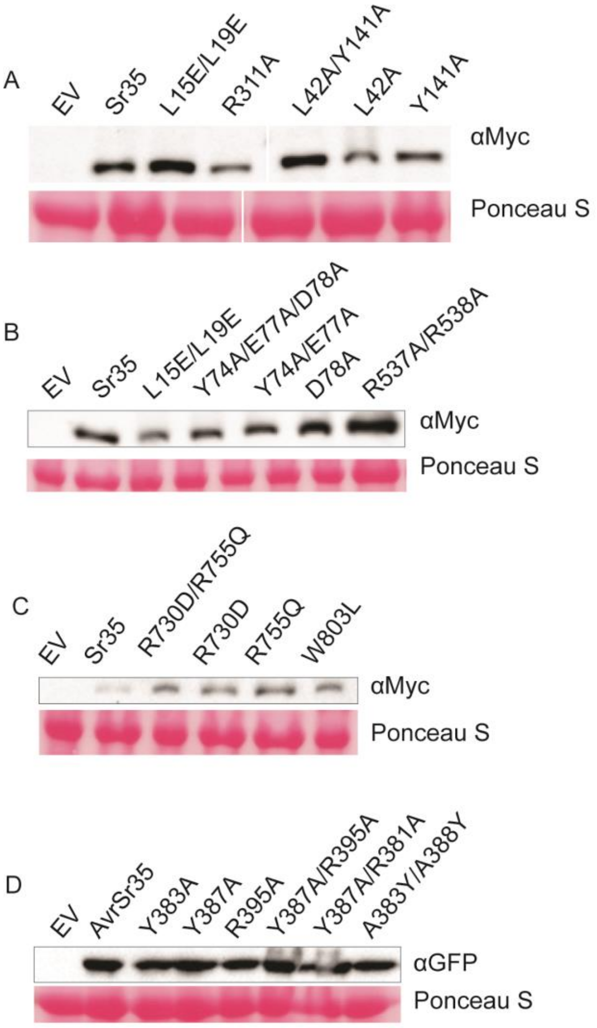

**Fig. S4** Western blot of *N. benthamiana* experiments. Pooled samples from 3 technical replicates. Ponceau S staining as a loading control. (**A**) Sr35 NBD ATP-binding and CC protomer interface mutants. Myc-tagged protein. (**B**) Sr35 EDVID and arginine-cluster mutants. Myc-tagged protein. (**C**) Sr35 LRR mutants. Myc-tagged protein. (**D**) AvrSr35 mutants. YFP-tagged protein detected by GFP antibody.

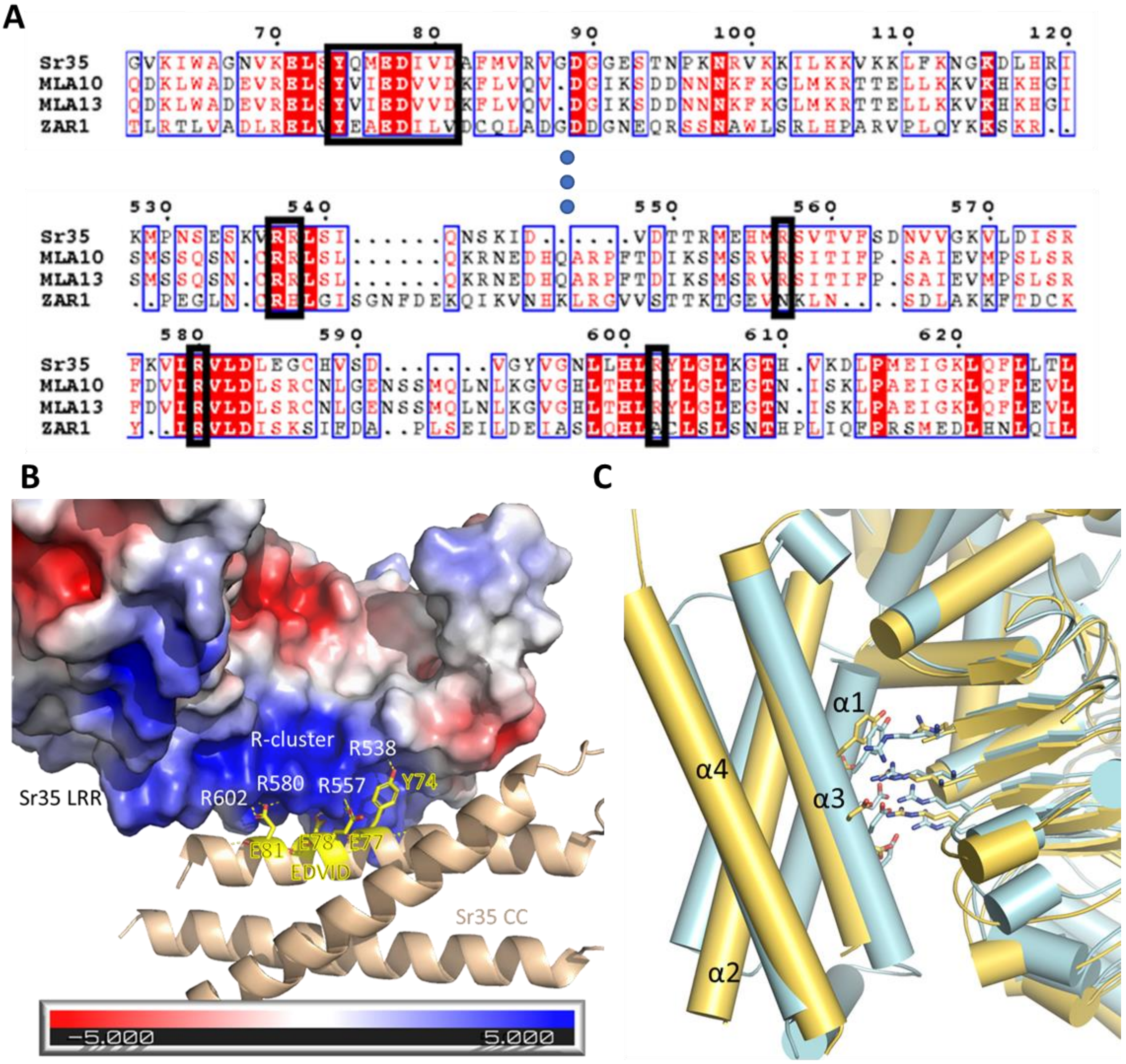


**Fig. S5** Details of EDIVD and R-cluster. (**A**) Multiple protein sequence alignment of *Hv*MLA10, *Hv*MLA13, Sr35 and ZAR1. Amino acids highlighted in red and in red text are identical and possess similar properties, respectively. Alignment of the EDIVD motif and arginine cluster are boxed in black (Robert and Gouet 2014). (**B**) Electrostatic surface charge of Sr35 LRR around the EDVID motif. (**C**) Structural alignment of Sr35 inactive structure prediction (cyan) and one protomer (yellow) from Sr35 resistosome. Detail view of EDVID and arginine cluster interactions. In analogy to ZAR1, the Sr35 CC α1-helix might undergo structural rearrangement, which likely requires EDVID with arginine cluster interactions to transiently resolve.

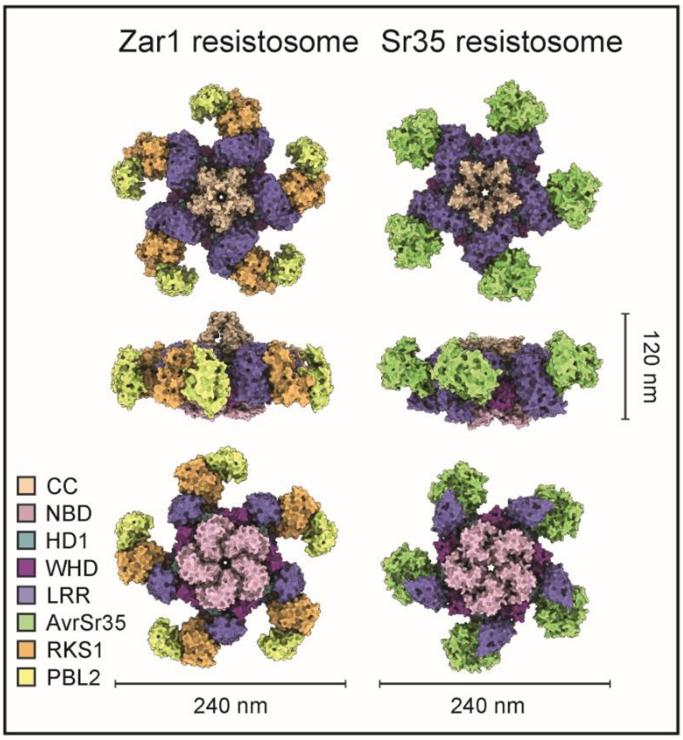


**Fig. S6** CNL resistosome structural conservation. The structures (in surface representation) of the ZAR1 resistosome and the Sr35 resistosome are shown. Zar1 is indirectly activated by the host proteins PBL2 and RKS1. Sr35 is directly activated by the fungal effector AvrSr35. The first, second, and third row show the top, side, and bottom views of these structures, respectively. Domains are coloured according to in-figure legend. Sizes are indicated by scale bar.

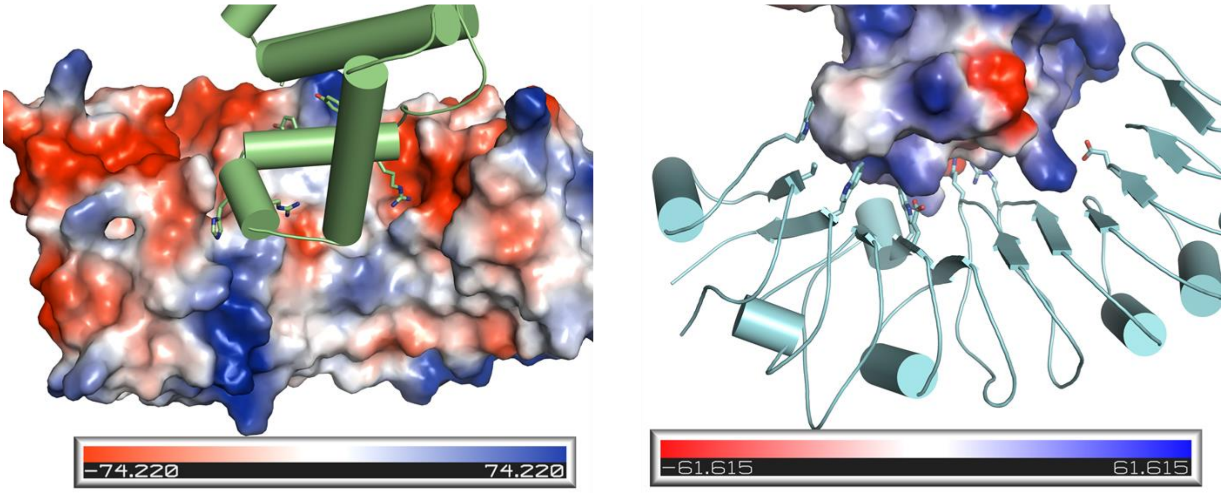


**Fig. S7** Shape and charge complementarity of Sr35 LRR and AvrSr35 at their interface. (Left) AvSr35 shown as cartoon (lime) and Sr35 as electrostatics surface model. (Right) Sr35 LRR shown as cartoon (cyan) and AvrSr35 as electrostatics surface model.

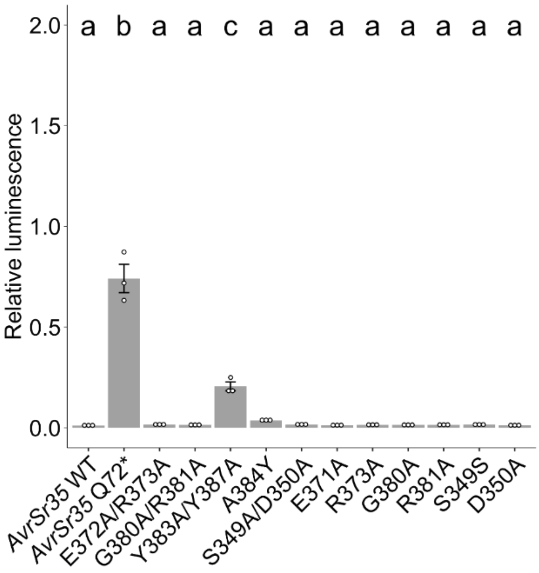


**Fig. S8** Wheat protoplast data of AvrSr35 mutants aiming to affect Sr35 recognition. Relative luminescence was used as a readout for cell death. N=3 replicates. Statistical test=one-way ANOVA, Tukeys test for multiple comparisons. Samples labelled with different letters differed significantly ($p < 0.05$) in the statistical test.

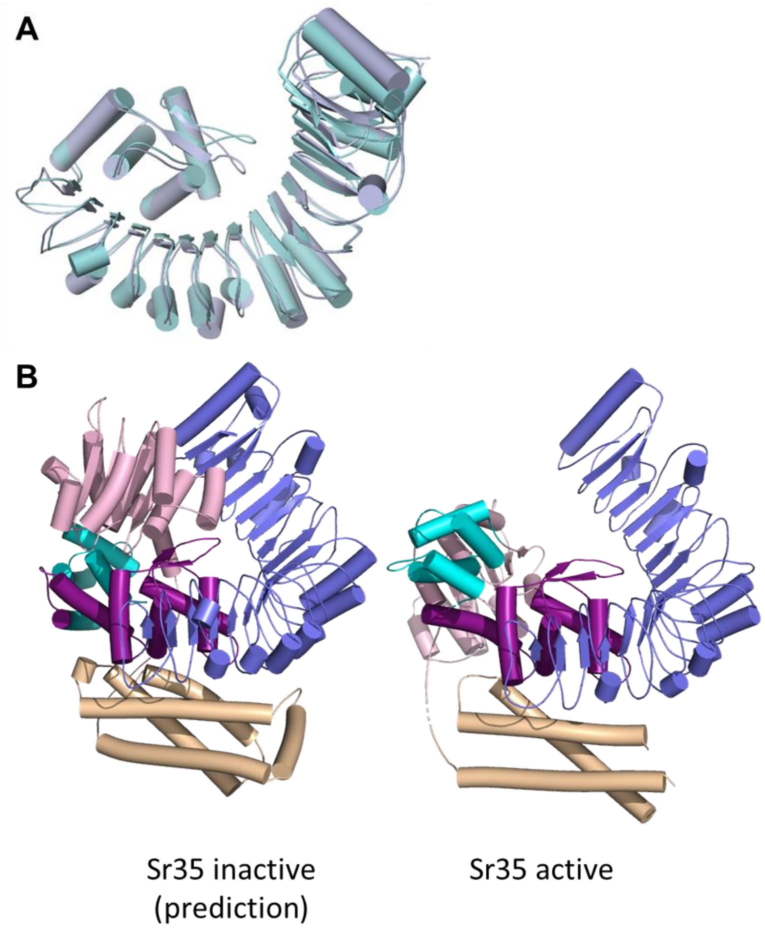


**Fig. S9** Comparison Sr35 prediction (Alphafold 2) and Sr35 protomer from Cryo-EM structure. (**A**) Structural alignment of WHD and LRR domains from Sr35 Alphafold 2 prediction (cyan) and from Sr35 resistosome Cryo EM structure (blue). (**B**) Structural comparison of monomeric Sr35 from prediction (left) and from Cryo-EM structure (right). Substantial differences exist highlighting the structural re-organization within the NOD module (NBD-HD1 relative to WHD). Domain color code: CC (yellow), NBD (light pink), HD1 (cyan), WHD (purple), and LRR (blue).

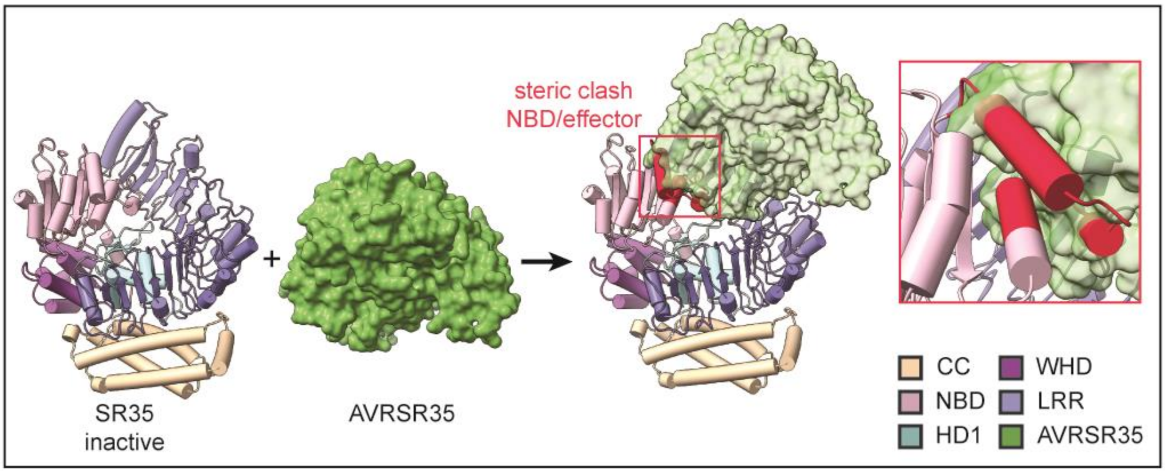


**Fig. S10** Steric clash between AvrSr35 and Sr35 NBD mediates Sr35 receptor activation**.** Inactive Sr35 inside the cell comes in contact with *Pgt* effector AvrSr35. In avoidance of a steric clash (red) between AvrSr35 and the Sr35 NBD domain, the Sr35 NBD domain is forced to structurally rearrange and an ‘primed’ receptor-effector complex is formed. Full activation and oligomerization requires subsequent ADP release, ATP binding and, NOD module rearrangement and CC domain structural rearrangement. Sr35 domains and AvrSr35 coloured according to in-figure legend.

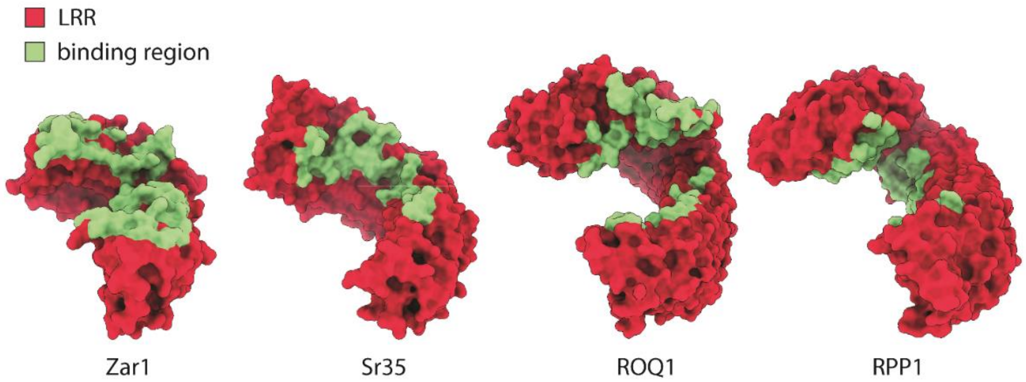


**Fig. S11** Comparison ZAR1, Sr35, ROQ1, RPP1 ligand binding site. Ligand binding to LRR of CNLs (Zar1, Sr35) and LRR-CJID of TNLs (Roq1, RPP1) occurs in analogous region in the ascending lateral LRR domain.

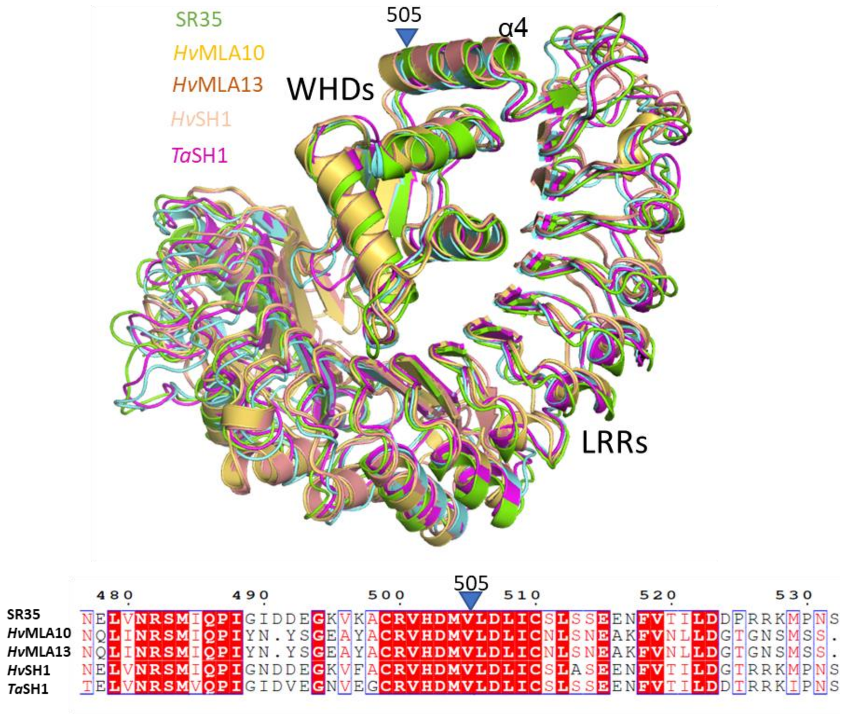


**Fig. S12** Rationale for chimera design; structural (top) and sequence (bottom) alignment of Sr35, *Hv*MLA10, *Hv*MLA13, *Ta*SH1 and *Hv*SH1. Amino acid 505 in the structurally and sequence conserved α4-helix of the WHD of Sr35 was included in hybrid CNL receptors.

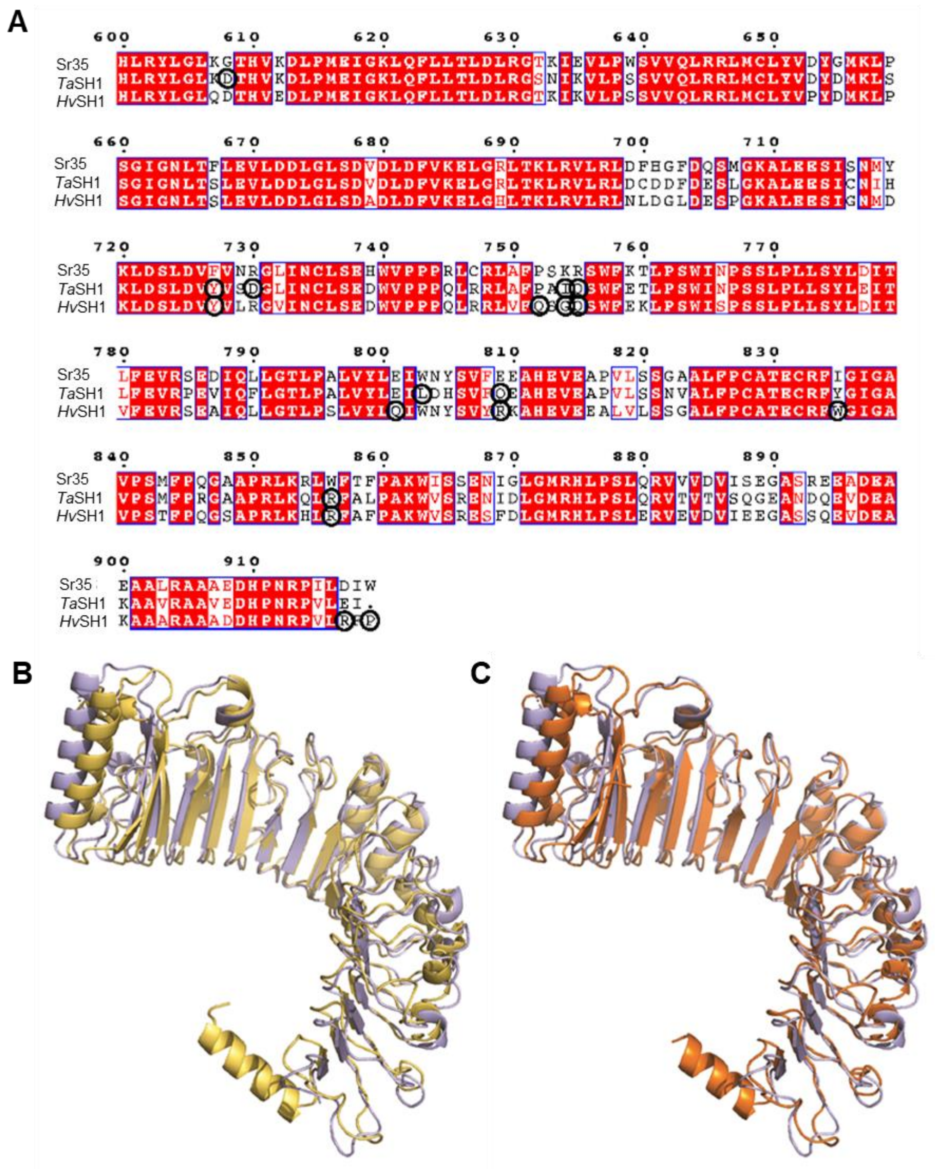


**Fig. S13** (**A**) Multiple protein sequence alignment of Sr35, TaSH1 and HvSH1. Circled amino acids were substituted to corresponding amino acids in the Sr35 sequence for the generation of *TaSh1*$^{GOF}$ and *HvSh1*$^{GOF}$ constructs. Amino acids highlighted in red and in red text are identical and possess similar properties, respectively (Robert and Gouet 2014). (**B**) Structural alignment of Sr35 LRR (light blue) with structural prediction of *Ta*SH1 (yellow) and (**C**) *Hv*SH1 (orange).
